# Placental effects on maternal brain revealed by disrupted placental gene expression in mouse hybrids

**DOI:** 10.1101/527143

**Authors:** Lena Arévalo, Polly Campbell

**Affiliations:** Department of Integrative Biology, Oklahoma State University, Stillwater, OK, U.S.A.

**Keywords:** Placenta, Maternal behavior, hybrid, Maternal-fetal communication, Maternal-fetal conflict, Imprinted genes, Placenta-specific gene families, Placental lactogens, Dopamine, Paternal effects, *Mus musculus domesticus*, *Mus spretus*, Genomic imprinting

## Abstract

The mammalian placenta is both the physical interface between mother and fetus, and the source of endocrine signals that target the maternal hypothalamus, priming females for parturition, lactation and motherhood. Despite the importance of this connection, the effects of altered placental signaling on the maternal brain are understudied. Here, we show that placental dysfunction alters gene expression in the maternal brain, with the potential to affect maternal behavior. Using a cross between the house mouse and the Algerian mouse in which hybrid placental development is abnormal, we sequenced late gestation placental and maternal medial preoptic area transcriptomes and quantified differential expression and placenta-maternal brain co-expression between normal and hybrid pregnancies. The expression of *Fmn1, Drd3, Caln1* and *Ctsr* was significantly altered in the brains of females exposed to hybrid placentas. Most strikingly, expression patterns of placenta-specific gene families and *Drd3* in the brains of house mouse females carrying hybrid litters matched those of female Algerian mice, the paternal species in the cross. Our results indicate that the paternally-derived placental genome can influence the expression of maternal-fetal communication genes, including placental hormones, suggesting an effect of the offspring’s father on the mother’s brain.

## Introduction

The placenta is a unique, transient organ shared by two organisms. Placental morphology is surprisingly diverse across vertebrates and is subject to rapid evolutionary change and convergent evolution (Blackburn 1993; Reznick et al. 2002; Roberts et al. 2016; Armstrong et al. 2017). In most eutherian mammals, including mice and humans, successful blastocyst implantation relies on endometrial invasion by the embryonic trophoblast cells that give rise to the mature placenta (Cross et al. 1994). As such, the placenta provides the closest physical and molecular link between mother and offspring seen in any animal (Wagner et al. 2014). This intimate connection promotes an array of maternal-fetal interactions, including bidirectional hormonal regulation and even the exchange of entire cells. These interactions are not spatially limited, but extend to both the fetal and the maternal brain (Bridges et al. 1996; Ladyman et al. 2010; Boddy et al.2015).

Throughout pregnancy the placenta mediates the regulation of resource allocation, immune tolerance, fetal development and, importantly, hormonal priming of the maternal brain. A key subset of placenta-secreted molecules are involved in priming maternal physiology for parturition and lactation, and promoting the onset of maternal behaviors in late gestation (reviewed in Bridges 2015; Creeth et al. 2018). In rodents, the medial preoptic area (MPoA) in the hypothalamus is thought to be the primary neural target of these placental molecules (Bridges et al. 1996; Mann and Bridges 2001; Larsen and Grattan 2012). The MPoA has been characterized as the central hub of parenting behavior (Kohl and Dulac 2018): receptors for key pregnancy hormones and neurotransmitters, including estrogen, prolactin and dopamine, are highly expressed in this nucleus and interact with ligands of both maternal and placental origin (Numan and Insel 2003).

Two classes of placental genes are of particular importance to the interaction between placenta and maternal brain: imprinted genes (IGs) and placenta-specific gene families (PSFs). Several IGs and PSF genes are expressed in the same placental compartment, especially in the placental endocrine compartment (Simmons et al. 2007, 2008; Tunster et al. 2013). IGs are exclusively or predominantly expressed from one allele, and are highly expressed in placenta and brain. The allelic expression is initiated by heritable epigenetic marks (“imprints”) in maternal and paternal germ cells, such that some IGs are maternally silenced and paternally expressed, whereas others are paternally silenced and maternally expressed (Reik and Walter 2001; Ferguson-Smith 2011). During pregnancy, IGs are critical to placental development and function, maintaining the balance between maternal supply and embryonic demand, and regulating maternal-fetal exchange (Constancia et al. 2005; Lefebvre 2012; Tunster et al. 2013).

PSFs arose through lineage-specific gene duplication events during placental evolution (Rawn and Cross 2008). In rodents, these are the prolactin gene family (placental lactogens (Prls)), placental cathepsin proteases and their inhibitors (PECs) and pregnancy specific glycoproteins (PSGs) (Zebhauser et al. 2005; Soares et al. 2007; Mason 2008). PSF gene products are mainly expressed from the placental endocrine compartment and many are secreted into the maternal bloodstream; key functions include placental development, immunoregulation, and physiological and neurological priming of the maternal organism (Rawn and Cross 2008; Bridges et al. 1996; Mann and Bridges 2001). Most notably, a subset of PRLs are able to bind the prolactin receptor that is expressed in the maternal MPoA leading to the proposal that placental hormones directly affect maternal endocrine state and behavior (Bridges et al. 1996; Larsen and Grattan 2012). IGs are implicated in regulating PSF secretion via their effects on the structure and function of the placental endocrine compartment (John 2017). However, our current understanding of the role of IGs in PSF signaling is rudimentary, and the relationship between gene expression in placenta and the maternal MPoA is uncharted.

The majority of the placenta, including the endocrine compartment, is fetally-derived. Placental representation of both parental genomes sets the stage for conflict (maternal-paternal and parent-offspring), and for coadaptation (mother-offspring), with imprinted genes uniquely positioned to mediate both types of interactions (Moore and Haig 1991; Wolf and Hager 2006, Keverne and Curly 2008; Haig 2014). However, while evolutionary models for imprinted gene expression abound (reviewed in Patten et al. 2014), few have considered the interaction between paternally-derived placental signals and signal reception in the maternal brain (Janssen et al. 2016a; McNamara et al. 2018; Creeth et al. 2018).

Here, we use a natural hybrid system to explore the effects of placental dysregulation on gene expression in the maternal brain. Over- or under-growth that depends on the direction of the cross is a signature of disrupted imprinting in mammalian hybrids (Vrana 2007). This pattern is documented in several orders (Gray 1972), with the best-studied examples in rodents (multiple species in the genera *Peromyscus, Mus* and *Phodopus* (Zechner et al. 1996; Vrana et al. 1998; Brekke and Good 2014)). Parent-of-origin growth effects in the cross between the house mouse, *Mus m. domesticus* (*Dom*) and the Algerian mouse, *M. spretus* (*Spret*), were first described over 20 years ago: placentas are undersized when the mother is *Dom* and the father is *Spret*, and severely oversized in the reciprocal cross, with more extreme size effects in both directions of the backcross (Zechner et al. 1996). Subsequent studies confirmed altered expression and methylation of candidate IGs, and disrupted placental organization (Hemberger et al. 1999; Zechner et al. 2002, 2004; Shi et al. 2005). Specifically, the placental endocrine compartment (or junctional zone) was shown to be reduced and disorganized (Zechner et al. 1996; Kurz et al. 1999). However, the extent of placental misexpression has not been measured on a genome scale, and this system’s potential to uncover the maternal consequences of altered placental signaling has not been considered.

The characteristics of this classic system, together with the availability of high quality genomes for both parental species (Keane et al. 2011), make the cross between *Dom* and *Spret* an excellent model in which to explore the effects of placental disruptions on the maternal brain. By comparing MPoA expression between females of the same species that differ only in the type of pregnancy/placenta they experience (hybrid vs. conspecific), we specifically isolate the effect of placental gene expression differences on the maternal brain (Fig. 1). Characterization of altered gene expression at the maternal-fetal interface provides insight into the mechanisms of maternal-fetal communication, the contribution of the paternal genome to this interaction, and identifies promising candidate genes for future evolutionary and biomedical work.

**Figure 1.**
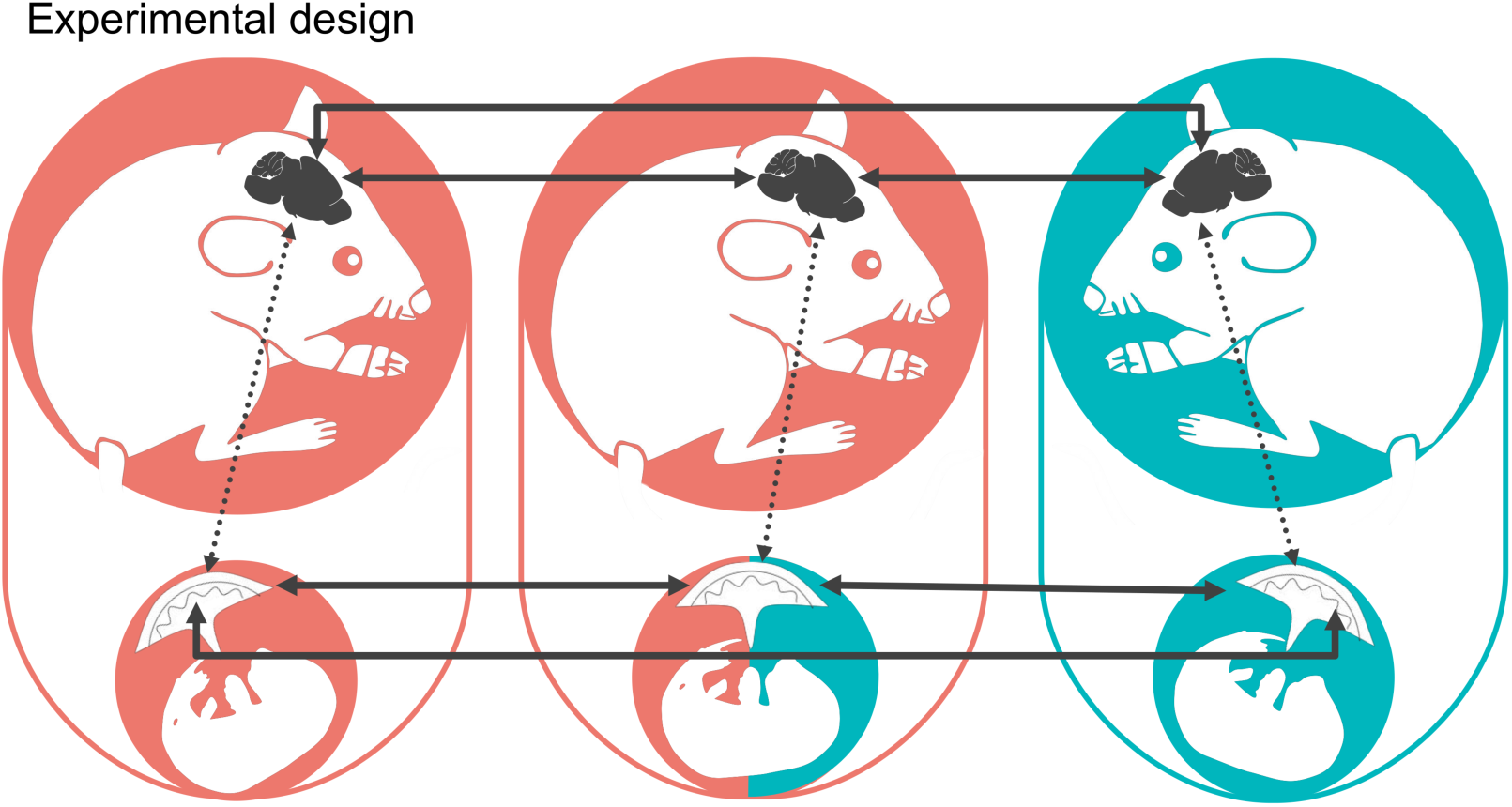
Experimental design. Schematic representation of the comparisons performed to test for differential gene expression in the medial preoptic area of the maternal brain and the placenta during late gestation. *Mus m. domesticus* is depicted in red and *Mus spretus* in blue. Hybrid tissue is indicated by a combination of red and blue. Solid arrows indicate differential gene expression analysis. Dashed arrows indicate co-expression analysis.

## Results

To study the relationship between placenta and maternal MPoA on a molecular level we produced three crosses resulting in three types of pregnancy: *Dom* x *Dom* (*Dom* pregnancy, n=5, litter size range = 4-6, sex ratio range [proportion males] = 0.3-0.6), *Dom* x *Spret* (hybrid pregnancy, n=5, litter size range = 3-6, sex ratio range = 0.3-1.0) and *Spret* x *Spret* (*Spret* pregnancy, n=5, litter size range = 5-6, sex ratio range = 0.2-0.8) (in all crosses, female is first). For each type of cross we produced 5 biological replicates and extracted the maternal MPoA and placentas from each pregnant female in late gestation at embryonic day (e) 17.5. During late gestation the effect of placental signaling on the maternal MPoA is specifically important for the onset of maternal care at parturition (Bridges et al. 1996; Mann and Bridges 2001; Larsen and Grattan 2012). Placenta weights were compared to confirm that hybrid placentas were hypoplastic as expected (Zechner et al. 1996). Since litter size can have an effect on placental and embryo size we corrected weights for litter size. Hybrid placentas weighed significantly less than placentas of both parental species with no significant differences between parental species (One-way ANOVA (Females): F(2)=56.95, p<0.001, Tukey HSD: Hybrid-Dom: diff=-0.1, p<0.001, Hybrid-Spret: diff=0.1, p<0.001, Dom-Spret: diff=0.002, p=0.97). We sequenced the maternal MPoA transcriptome and the placental transcriptomes of 1 male and 1 female per mother (n=9-10/type of pregnancy, in hybrid pregnancies one litter consisted of only males, thus n=4 for females), and evaluated differential expression between all pregnancy types (Fig. 1). Because the maternal brain is exposed to the placental signals of both sexes simultaneously, male and female placental expression was analyzed jointly. Additionally, we assessed co-expression between the two tissues for each type of pregnancy and determined the differences in co-expression between pregnancy types (Fig. 1).

### Differential expression in the placenta

We tested for differential expression in three pairwise comparisons: hybrid vs. *Dom*, hybrid vs. *Spret*, and *Dom* vs. *Spret* placentas. For placental comparisons, only genes with log2 fold change (LFC) in expression ≥ 0.5 (1.5 times higher or lower expression), and Benjamini-Hochberg-corrected p≤0.05, were considered significantly differentially expressed (DE). We define transgressive expression in hybrids as expression that is significantly higher or lower compared to both parental species. Hybrid genes that were DE compared to *Dom* but not to *Spret* are defined as having *Spret*-like expression patterns, and vice versa.

In hybrid placentas 9.73% of all tested genes were expressed higher and 7.79% lower compared to *Dom* placentas (up: 1,781/18,298, including 11 IGs; down: 1,426/18,298, including 3 IGs) (Fig. 2, Supplemental Fig. S1 and Supplemental Dataset S1). Compared to *Spret* placentas, 16.32% of genes were expressed higher and 9.6% lower in hybrids (up: 3,036/18,529, including 16 IGs; down: 1,801/18,529, including 7 IGs) (Fig. 2, Supplemental Fig. S2, Supplemental Dataset S1). Thirty-two percent of all tested genes were DE between *Dom* and *Spret* (up: 4,014/19,079, including 19 IGs; down: 3,278/19,079, including 13 IGs) (Supplemental Fig. S3, Supplemental Dataset S1). Hybrids clustered slightly closer to Dom samples in the diagnostic PCA plot (Supplementary Fig. 4).

**Figure 2.**
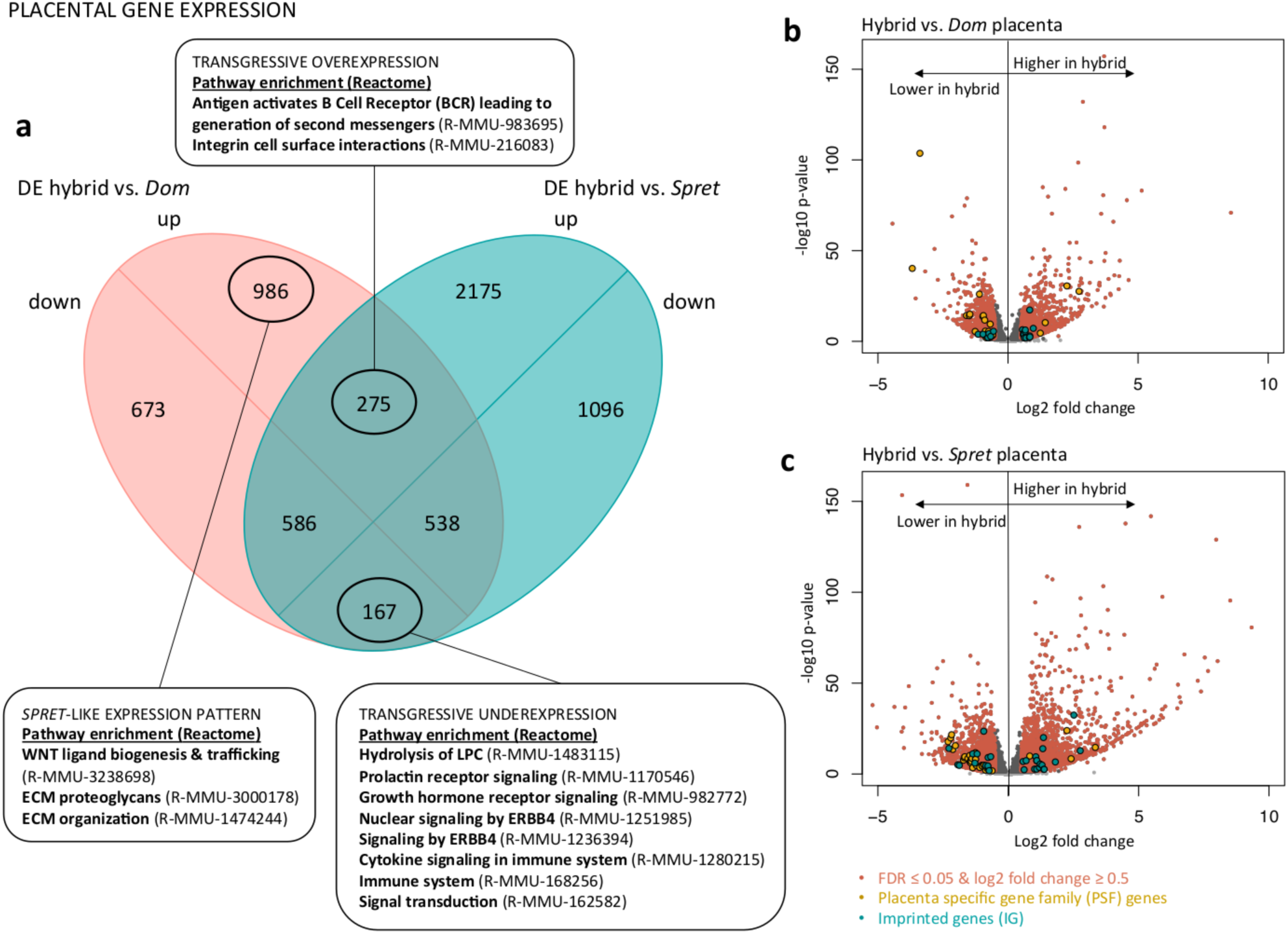
Placental gene expression. Summary of results of differential gene expression (DE) analysis between *Mus m. domesticus* (*Dom*), *Mus spretus* (*Spret*) and hybrid placentas. (A) Venn diagram indicating the overlap of differentially expressed genes between the comparisons hybrid vs. *Dom* and hybrid vs. *Spret*; up = genes expressed higher in hybrids compared to parental species, down = genes expressed lower in hybrids compared to parental species. Genes expressed higher or lower compared to both parental species (transgressive expression) and genes with *Spret*-like expression in the hybrid are marked in the diagram. Results of pathway overrepresentation (Reactome, version 58,Mi et al. 2017) are provided in connected text boxes. Volcano plot of DE analysis results of (B) hybrid vs. *Dom* and (C) hybrid vs. *Spret* placentas. Significantly differentially expressed genes with FDR ≤ 0.05 and log2 fold change ≥ 0.5 are depicted in red. Imprinted genes are indicated in blue and placenta-specific gene family genes in yellow.

To explain hybrid placental phenotypes that are not intermediate to both parents, genes with transgressive expression are of specific interest. We found 275 genes that were expressed at higher, and 167 at lower, levels in hybrids compared to both parental species (Fig. 2*A*). Transgressively higher expressed genes were significantly enriched for B-cell receptor activation and integrin cell surface interaction pathways (Fig. 2*S*, Supplemental Dataset S1). Interestingly, transgressively lower expressed genes were enriched for prolactin and growth hormone receptor signaling, ERBB signaling, and cytokine signaling in immune system, among others (Fig. 2*A*, Supplemental Dataset S1). Many Prls are involved in these pathways, along with other genes. PSF genes were highly overrepresented among DE genes in hybrids compared to both *Dom* (Fisher’s exact test: p<0.001, odds ratio = 5.65) and *Spret* (p<0.001, odds ratio = 3.41). Multiple members of these gene families were misexpressed in the hybrid placenta, with the majority being expressed lower compared to both parental species (14 transgressively lower, 7 DE intermediate) (Table 1).

**Table 1.**
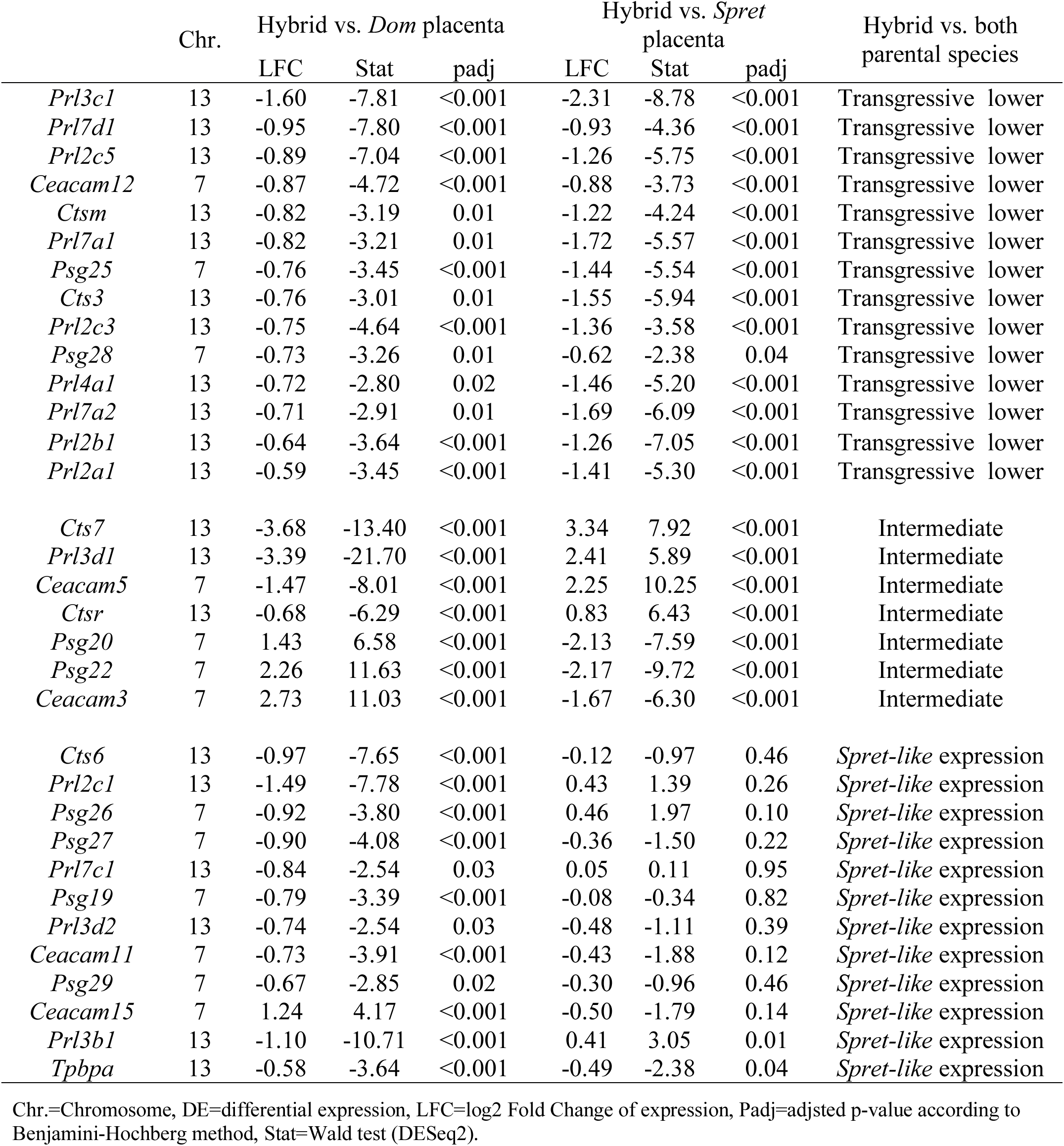
Differential expression of PSF genes in hybrid placenta compared to both parental species

|  | Chr. | Hybrid vs. *Dom* placenta |  |  | Hybrid vs. *Spret* placenta |  |  | Hybrid vs. both parental species |
| --- | --- | --- | --- | --- | --- | --- | --- | --- |
|  |  | LFC | Stat | padj | LFC | Stat | padj |  |
| *Prl3c1* | 13 | -1.60 | -7.81 | <0.001 | -2.31 | -8.78 | <0.001 | Transgressive lower |
| *Prl7d1* | 13 | -0.95 | -7.80 | <0.001 | -0.93 | -4.36 | <0.001 | Transgressive lower |
| *Prl2c5* | 13 | -0.89 | -7.04 | <0.001 | -1.26 | -5.75 | <0.001 | Transgressive lower |
| *Ceacam12* | 7 | -0.87 | -4.72 | <0.001 | -0.88 | -3.73 | <0.001 | Transgressive lower |
| *Ctsm* | 13 | -0.82 | -3.19 | 0.01 | -1.22 | -4.24 | <0.001 | Transgressive lower |
| *Prl7a1* | 13 | -0.82 | -3.21 | 0.01 | -1.72 | -5.57 | <0.001 | Transgressive lower |
| *Psg25* | 7 | -0.76 | -3.45 | <0.001 | -1.44 | -5.54 | <0.001 | Transgressive lower |
| *Cts3* | 13 | -0.76 | -3.01 | 0.01 | -1.55 | -5.94 | <0.001 | Transgressive lower |
| *Prl2c3* | 13 | -0.75 | -4.64 | <0.001 | -1.36 | -3.58 | <0.001 | Transgressive lower |
| *Psg28* | 7 | -0.73 | -3.26 | 0.01 | -0.62 | -2.38 | 0.04 | Transgressive lower |
| *Prl4a1* | 13 | -0.72 | -2.80 | 0.02 | -1.46 | -5.20 | <0.001 | Transgressive lower |
| *Prl7a2* | 13 | -0.71 | -2.91 | 0.01 | -1.69 | -6.09 | <0.001 | Transgressive lower |
| *Prl2b1* | 13 | -0.64 | -3.64 | <0.001 | -1.26 | -7.05 | <0.001 | Transgressive lower |
| *Prl2a1* | 13 | -0.59 | -3.45 | <0.001 | -1.41 | -5.30 | <0.001 | Transgressive lower |
| *Cts7* | 13 | -3.68 | -13.40 | <0.001 | 3.34 | 7.92 | <0.001 | Intermediate |
| *Prl3d1* | 13 | -3.39 | -21.70 | <0.001 | 2.41 | 5.89 | <0.001 | Intermediate |
| *Ceacam5* | 7 | -1.47 | -8.01 | <0.001 | 2.25 | 10.25 | <0.001 | Intermediate |
| *Ctsr* | 13 | -0.68 | -6.29 | <0.001 | 0.83 | 6.43 | <0.001 | Intermediate |
| *Psg20* | 7 | 1.43 | 6.58 | <0.001 | -2.13 | -7.59 | <0.001 | Intermediate |
| *Psg22* | 7 | 2.26 | 11.63 | <0.001 | -2.17 | -9.72 | <0.001 | Intermediate |
| *Ceacam3* | 7 | 2.73 | 11.03 | <0.001 | -1.67 | -6.30 | <0.001 | Intermediate |
| *Cts6* | 13 | -0.97 | -7.65 | <0.001 | -0.12 | -0.97 | 0.46 | *Spret-like* expression |
| *Prl2c1* | 13 | -1.49 | -7.78 | <0.001 | 0.43 | 1.39 | 0.26 | *Spret-like* expression |
| *Psg26* | 7 | -0.92 | -3.80 | <0.001 | 0.46 | 1.97 | 0.10 | *Spret-like* expression |
| *Psg27* | 7 | -0.90 | -4.08 | <0.001 | -0.36 | -1.50 | 0.22 | *Spret-like* expression |
| *Prl7c1* | 13 | -0.84 | -2.54 | 0.03 | 0.05 | 0.11 | 0.95 | *Spret-like* expression |
| *Psg19* | 7 | -0.79 | -3.39 | <0.001 | -0.08 | -0.34 | 0.82 | *Spret-like* expression |
| *Prl3d2* | 13 | -0.74 | -2.54 | 0.03 | -0.48 | -1.11 | 0.39 | *Spret-like* expression |
| *Ceacam11* | 7 | -0.73 | -3.91 | <0.001 | -0.43 | -1.88 | 0.12 | *Spret-like* expression |
| *Psg29* | 7 | -0.67 | -2.85 | 0.02 | -0.30 | -0.96 | 0.46 | *Spret-like* expression |
| *Ceacam15* | 7 | 1.24 | 4.17 | <0.001 | -0.50 | -1.79 | 0.14 | *Spret-like* expression |
| *Prl3b1* | 13 | -1.10 | -10.71 | <0.001 | 0.41 | 3.05 | 0.01 | *Spret-like* expression |
| *Tpbpa* | 13 | -0.58 | -3.64 | <0.001 | -0.49 | -2.38 | 0.04 | *Spret-like* expression |

Chr.=Chromosome, DE=differential expression, LFC=log2 Fold Change of expression, Padj=adjusted p-value according to Benjamini-Hochberg method, Stat=Wald test (DESeq2).

Notably, approximately twice as many genes in hybrid placentas were uniquely DE relative to *Spret* (3,271) as opposed to *Dom* (1,659) (Fig. 2*A*). Thus, the general expression pattern in hybrid placentas was more similar to the maternal species. *Dom*-like expressed genes were enriched for multiple immune related pathways, together with angiogenesis, vascular development and hemostasis related terms, among others (Supplemental Dataset S1).

Genes with a *Spret*-like expression pattern in hybrid placentas are of particular interest, since these have the potential to alter communication with the *Dom* maternal brain. Of the 1,659 genes with *Spret-*like expression (Fig. 2*A*), 12 were PSF genes and 6 were IGs (Tables 1 and 2). *Spret*-like expressed genes in the hybrid were enriched for WNT signaling and extracellular matrix organization pathways, among others (Fig. 2*A*, Supplemental Dataset S1).

**Table 2.** Differential expression of imprinted genes in hybrid placenta compared to both parental species

|  | IC/Chr. | Hybrid vs. *Dom*<br>placenta |  |  | Hybrid vs. *Spret* placenta |  |  | Hybrid vs. both parental<br>species | expressed<br>allele |
| --- | --- | --- | --- | --- | --- | --- | --- | --- | --- |
|  |  | LFC | Stat | padj | LFC | Stat | padj |  |  |
| *Phlda2* | dist7-IC2 | 0.97 | 5.39 | <0.001 | 1.09 | 5.44 | <0.001 | Transgressive higher | Maternal |
| *Klf14* | prox6 | 0.67 | 5.01 | <0.001 | 1.06 | 6.29 | <0.001 | Transgressive higher | Maternal |
| *Tnfrsf23* | dist7-IC2 | 0.59 | 3.57 | <0.001 | 1.97 | 9.02 | <0.001 | Transgressive higher | Maternal |
| *Tspan32* | dist7-IC2 | -0.66 | -3.11 | 0.01 | 2.01 | 6.43 | <0.001 | Intermediate | Maternal |
| *Th* | dist7 | 0.69 | 2.48 | 0.04 | -2.28 | -7.76 | <0.001 | Intermediate | Maternal |
| *Ascl2* | dist7-IC2 | -0.96 | -3.85 | <0.001 | -1.28 | -4.90 | <0.001 | Transgressive lower | Maternal |
| *Sfmbt2* | 2 | -0.56 | -4.69 | <0.001 | -0.67 | -6.30 | <0.001 | Transgressive lower | Paternal |
| *Magel2* | cent7 | 0.61 | 3.81 | <0.001 | 0.48 | 2.25 | 0.06 | *Spret-like* expression | Paternal |
| *Dcn* | 10 | 0.63 | 2.45 | 0.04 | -0.06 | -0.16 | 0.92 | *Spret-like* expression | Maternal |
| *Nap1l5* | prox6 | 0.82 | 2.63 | 0.03 | 0.31 | 0.91 | 0.49 | *Spret-like* expression | Paternal |
| *Grb10* | prox11 | 0.83 | 8.65 | <0.001 | -0.06 | -0.47 | 0.74 | *Spret-like* expression | Maternal |
| *Nnat* | dist2 | 0.84 | 3.29 | <0.001 | -0.07 | -0.27 | 0.86 | *Spret-like* expression | Paternal |
| *Igf2* | dist7-IC1 | 0.84 | 2.89 | 0.01 | 0.02 | 0.08 | 0.96 | *Spret-like* expression | Paternal |

IC=imprinting cluster (https://www.mousebook.org (03.22.18)), Chr.=Chromosome, DE=differential expression, LFC=log2 Fold Change of expression, Padj=adjusted p-value according to Benjamini-Hochberg method, Stat=Wald test (DESeq2), expressed allele according to https://www.mousebook.org (03.22.18).

IGs were significantly overrepresented among hybrid DE genes compared to both *Spret* (Fisher’s exact test: p=0.02, odds ratio=0.59) and *Dom* (p=0.03, odds ratio=0.55). Three IGs (*Tnfrsf23, Phlda2* and *Klf14*) were transgressively higher expressed and two, (*Ascl2* and *Sfmbt2*) were transgressively down-regulated. Two additional IGs, *Tspan32* and *Th*, were significantly DE compared to both parental species but intermediate between the two. Four of these misexpressed IGs belong to the same imprinting cluster (IC2) on the distal part of mouse chromosome 7 (dist7) (MouseBook (https://www.mousebook.org (03.22.18)), and are normally maternally expressed (Table 2, Supplemental Dataset S1). We note that *Tnfrsf23* is expressed in decidual cells at the junction between the maternally derived decidua and the extraembryonic placenta (Clark et al. 2002).

### Overlap of placental DE genes with genes involved in preeclampsia

Among the misexpressed genes in the hybrid placenta we noticed several that are also misexpressed in the human pregnancy pathology preeclampsia. Although mice do not develop preeclampsia, preeclampsia-like phenotypes can occur and several rodent models have been developed to study key symptoms such as hypertension, proteinura, and altered inflammatory response (Podjarny et al. 2004; Dokras et al. 2006). To test if DE genes in the hybrid overlap with genes related to preeclampsia we extracted genes associated with preeclampsia from the database for preeclampsia (http://ptbdb.cs.brown.edu/dbpec/ (Uzun et al. 2016)) and obtained the mouse orthologs for these from biomart (R-package biomaRt (Durinck et al. 2005)). Of the 490 mouse orthologs we obtained, 26 were transgressively misexpressed in the hybrid placenta. Preeclampsia related genes were significantly overrepresented among transgressively misexpressed genes (Fisher’s exact test: p=0.003, odds ratio=1.91). An altered inflammatory response at the fetal-maternal interface is involved in preeclampsia in humans (Harmon et al. 2016); in the hybrid placenta, both up- and down-regulated transgressively expressed genes were enriched for immune-related and cytokine signaling pathways (Supplemental Dataset S1).

### Differential expression in the MPoA

To explore maternal gene expression in response to placental genotype we compared late gestation MPoA of *Dom* females during *Dom* pregnancies (MPoA-*dom*), *Dom* females during hybrid pregnancies (MPoA-*hy*) and *Spret* females during *Spret* pregnancies (MPoA-*spret*). Neural and placental tissues were collected from the same females. Gene expression in MPoA-*hy* vs. MPoA-*dom* is expected to be highly similar, with any DE attributable to carrying a hybrid litter, while the other comparisons should result in a large number of DE genes attributable to interspecific differences. For the intraspecific comparison we report significantly DE genes with log2 fold change (LFC) ≥ 0.2, since differences are expected to be subtle. For all other comparisons we report significantly DE genes with LFC ≥ 0.5.

Four genes were DE between MPoA-*hy* and MPoA-*dom*: Cathepsin-R (*Ctsr*) was expressed higher in MPoA-*hy* compared to MPoA-*dom*, and Dopamine receptor 3 (*Drd3*), Calneuron 1 (*Caln1*) and Formin 1 (*Fmn1*) were expressed lower (up: 1/18,779; down: 3/18,779) (Fig. 3, Table 3, Supplemental Fig. S4 and Supplemental Dataset S2). In MPoA*-hy* compared to MPoA-*spret* 10.61% of all tested genes were expressed higher and 10.32% lower (up: 1,608/15,154; down: 1,565/15,154) (Fig. 3, Supplemental Fig. S5, Supplemental Dataset S2). In the interspecific comparison of conspecific pregnancies, 11.32% of all tested gene were expressed higher and 10.76% lower in MPoA*-dom* compared to MPoA-*spret* (up: 1,757/15,515; down: 1,670/15,515) (Fig. 3, Supplemental Fig. S6, Supplemental Dataset S2).

**Figure 3.**
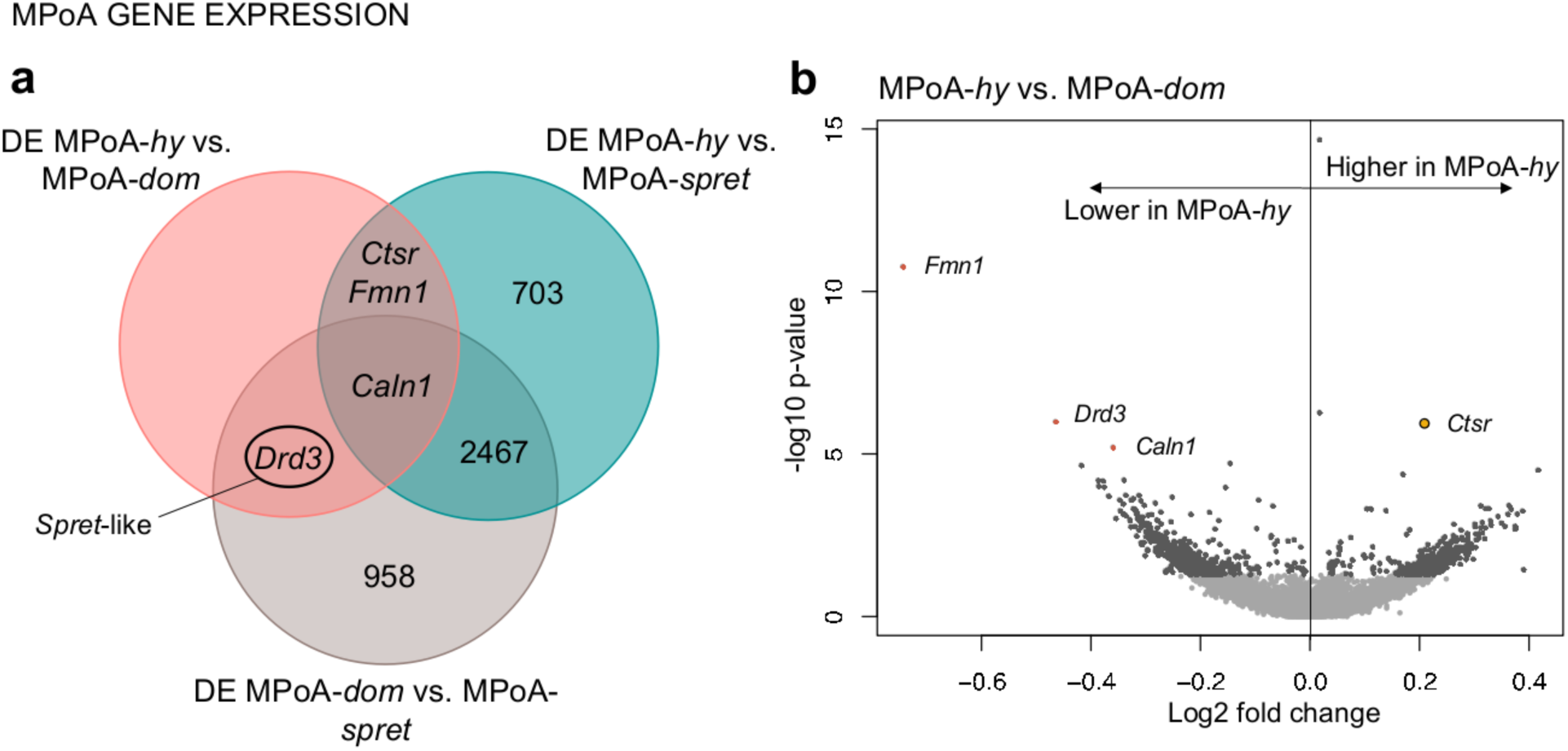
Maternal medial preoptic area (MPoA) gene expression. Summary of results of differential gene expression (DE) analysis between *Mus m. domesticus* MPoA during normal pregnancy (MPoA-*dom*), *Mus m. domesticus* MPoA during hybrid pregnancy (MPoA-*hy*) and *Mus spretus* MPoA during normal pregnacy (MPoA-*spret*). (A) Venn diagram indicating the overlap of differentially expressed genes between the comparisons MPoA-*hy* vs. MPoA-*dom*, MPoA-*hy* vs. MPoA-*spret* and MPoA-*dom* vs. MPoA-*spret*. Genes with *Spret*-like expression in MPoA-*hy* are marked in the diagram. (B) Volcano plot of DE analysis results of MPoA-*hy* vs. MPoA-*dom*. Significantly differentially expressed genes with FDR ≤ 0.05 and log2 fold change ≥ 0.2 are depicted in red, placenta-specific gene family genes in yellow.

**Table 3.** Differential expression in MpoA-*hy* compared to MPoA-*dom*.

|  | Chr. | LFC | Stat | padj |
| --- | --- | --- | --- | --- |
| *Fmn1* | 2 | -0.74 | -6.73 | <0.001 |
| *Drd3* | 16 | -0.47 | -4.89 | <0.001 |
| *Caln1* | 5 | -0.36 | -4.52 | 0.02 |
| *Ctsr* | 13 | 0.21 | 4.86 | <0.001 |

Chr.=Chromosome, LFC=log2 Fold Change of expression, Padj=adjusted p-value according to Benjamini-Hochberg method, Stat=Wald test (DESeq2).

Hybrid placentas express both *Spret* and *Dom* alleles and 1,659 genes had *Spret*-like expression. Therefore, the MPoA in *Dom* females carrying hybrid litters might exhibit expression patterns more similar to *Spret* female MPoA for some maternal-fetal communication genes. We extracted a list of 959 genes that were DE in MPoA-*dom* vs. MPoA-*spret* but not in MPoA-*hy* vs. MPoA-*spret*. This list includes *Drd3*, which was expressed lower in MPoA-*hy* compared to MPoA-*dom* (Fig. 3, Supplemental Dataset S2). Thus, *Drd3* could be defined as having *Spret*-like expression in MPoA-*hy. Fmn1* and *Ctsr* were expressed lower and higher, respectively, in MPoA-*hy* compared to both MPoA-*dom* and MPoA-*spret*, and were not DE between *Dom* and *Spret* MPoA during regular pregnancies (Fig. 3, Supplemental Dataset S2).

### Co-expression between the placenta and MPoA

The MPoA is thought to be an important target of placenta-secreted molecules and we found the placenta-specific gene, *Ctsr*, to be expressed in the maternal MPoA during hybrid pregnancies. Therefore, co-expression between placenta and MPoA is of particular interest. We determined the level of placenta-MPoA co-expression for the three pregnancy types (hybrid, *Dom*, *Spret*), and assessed differences between them.

10,839 genes were co-expressed between placenta and MPoA in all 3 comparisons. 172 genes were only co-expressed in *Dom* and 494 only in *Spret* pregnancies (Fig. 4, Supplemental Dataset S3). 176 genes were uniquely co-expressed in hybrid pregnancies. This gene set is of specific interest since these genes are indicators of MPoA response to abnormal placental expression. Uniquely co-expressed genes in hybrid pregnancies included 45 genes that were DE between hybrid and *Dom* placentas and 16 that were DE compared to both parental species’ placentas. *Ctsr*, which was significantly higher expressed in MPoA-*hy* compared to MPoA-*dom*, was uniquely co-expressed in hybrid pregnancies (Fig. 4, Supplemental Dataset S3).

**Figure 4.**
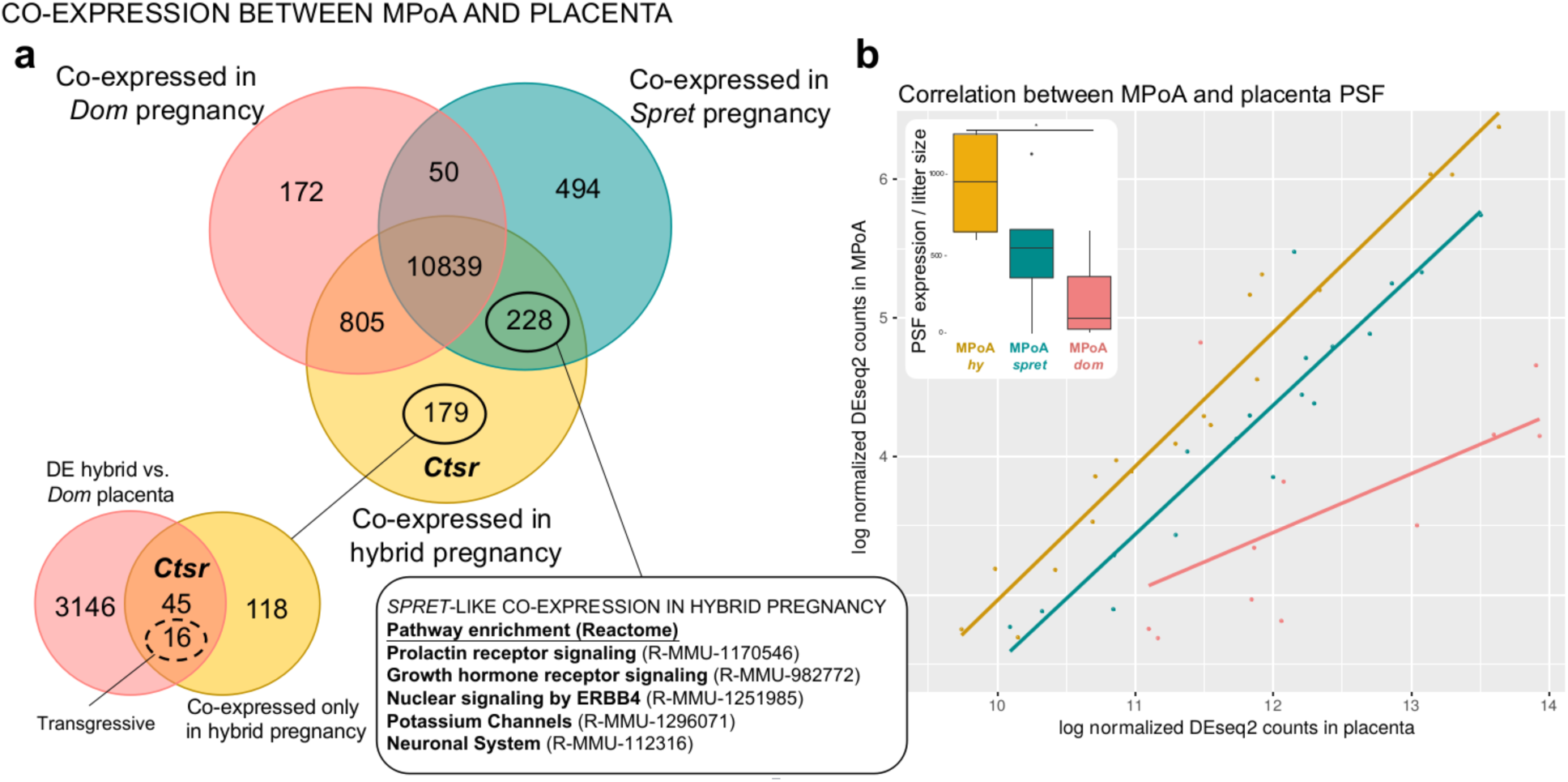
Co-expression between maternal medial preoptic area (MPoA) and placenta during late gestation. Summary of results of co-expression analysis for *Mus m. domesticus* during normal pregnancy (*Dom*-pregnancy), *Mus m. domesticus* during hybrid pregnancy (hybrid pregnancy) and *Mus spretus* during normal pregnancy (*Spret* pregnancy). (A) Venn diagram indicating the overlap of co-expressed genes between the three pregnancy types. Genes that are only co-expressed in hybrid pregnancy are marked in the diagram. A secondary Venn diagram for this gene set shows its overlap with differentially expressed genes in hybrid vs. *Dom* placentas. Transgressively expressed genes contained in this overlap are marked. Genes that are co-expressed in *Spret* and hybrid pregnancies but not in *Dom* pregnancies (*Spret*-like co-expression) are marked in the primary Venn diagram. Results of pathway overrepresentation (Reactome, version 58, Mi et al. 2017) for this gene set are provided in the connected text box. (B) Scatterplot showing the correlation between placenta specific gene family (PSF) gene expression in placenta and MPoA for the three pregnancy types. Red = *Dom* pregnancy (R^2^_adj_=0.38, p=0.01), Blue = *Spret* pregnancy (R^2^_adj_=0.9, p<0.001), Yellow = *hybrid* pregnancy (R^2^_adj_=0.95, p<0.001). Inset boxplot shows total PSF gene expression in MPoA (sum of normalized PSF counts/litter size), asterix indicates significant difference between MPoA-*hy* and MPoA-*dom*.

228 genes were co-expressed in *Spret* and hybrid pregnancies but not in *Dom* pregnancies. Thus, these genes exhibit *Spret*-like co-expression in the MPoA of *Dom* females carrying a hybrid litter. There was pathway overrepresentation overlap between this gene set and genes with transgressive misexpression in hybrid placenta (Fig. 2*A*), including prolactin and growth hormone receptor signaling, and ERBB signaling (Fig. 4, Supplemental Dataset S3). Moreover, PSFs were significantly overrepresented among these co-expressed genes (Fisher’s exact test: p<0.001, odds ratio=12.57). Although expression levels were far lower in the MPoA (range=10-589, mean=100 normalized counts, LFC to placenta: range=-9.54 to −14.13, mean=-11.12) than in the placenta (range=20-1,127,603, mean=97,988 normalized counts), these results are striking. To explore this relationship further, we tested for correlated expression of PSF genes between placenta and MPoA. We found very strong, positive correlations for hybrid (R^2^adj=0.95, p<0.001) and *Spret* (R^2^adj=0.9, p<0.001) and a significant but, surprisingly, weaker positive correlation for *Dom* (R^2^adj=0.38, p=0.01) (Fig. 4*B*, Table 3). Additionally, we found that total PSF expression (sum of all PSF read counts divided by litter size) is significantly higher in MPoA-*hy* relative to MPoA-*dom* but is statistically indistinguishable fromMPoA-*spret* (One-way ANOVA: F_2,12_=5.44, p=0.02, Tukey HSD: MPoA-*hy* vs. *MPoA-dom*: p=0.016, MPoA-*hy* vs. MPoA*-spret*: p=0.18, MPoA-*dom* vs. MPoA*-spret*: p=0.36) (Fig. 4*B*).

### Pairwise evolutionary rates of selected PSF genes

PSF genes were previously shown to exhibit accelerated evolutionary rates, potentially driven by maternal-fetal conflict (Chuong et al. 2010). We selected the top 10 co-expressed PSF genes with the highest expression in MPoA and extracted pairwise evolutionary rates (dN/dS) from Biomart (R-package biomaRt, (Durinck et al. 2005)) to test for evidence of positive selection. We also included the PSF gene *Ctsr*, which was significantly differentially expressed between MPoA-*hy* and MPoA-*dom*.

dN/dS, the per site ratio of nucleotide substitutions that change amino acid identity to those that do not, is an indicator of selective pressure, with dN/dS = 1 indicating neutral evolution, dN/dS < 1 purifying selection, and dN/dS > 1 diversifying positive selection (Goldman and Yang 1994). Of the 11 genes, three (*Prl8a6*, *Tpbpb* and *Ctsr*) had pairwise dN/dS >1 between *Dom* and *Spret* (Fig. 5, Supplemental Table S1). To infer which lineage experienced selection, we analyzed these three genes using the application CodeML (implemented in PAML 4; (Yang and Rannala 1997; Yang 2007)), and including sequences from additional *Mus* subspecies and species (*Mus m. musculus* (*Musc*), *Mus m. castaneus* (*Cast*), *Mus caroli* (*Car*) and *Mus pahari* (*Pah*)). CodeML fits alternative models to the data; the best-fit model is chosen based on likelihood ratio tests (LTRs). The two main models tested were M0, one evolutionary rate for the whole tree, and MC, selected branches (foreground) evolve at a different rate than the rest of the tree (background).

**Figure 5.**
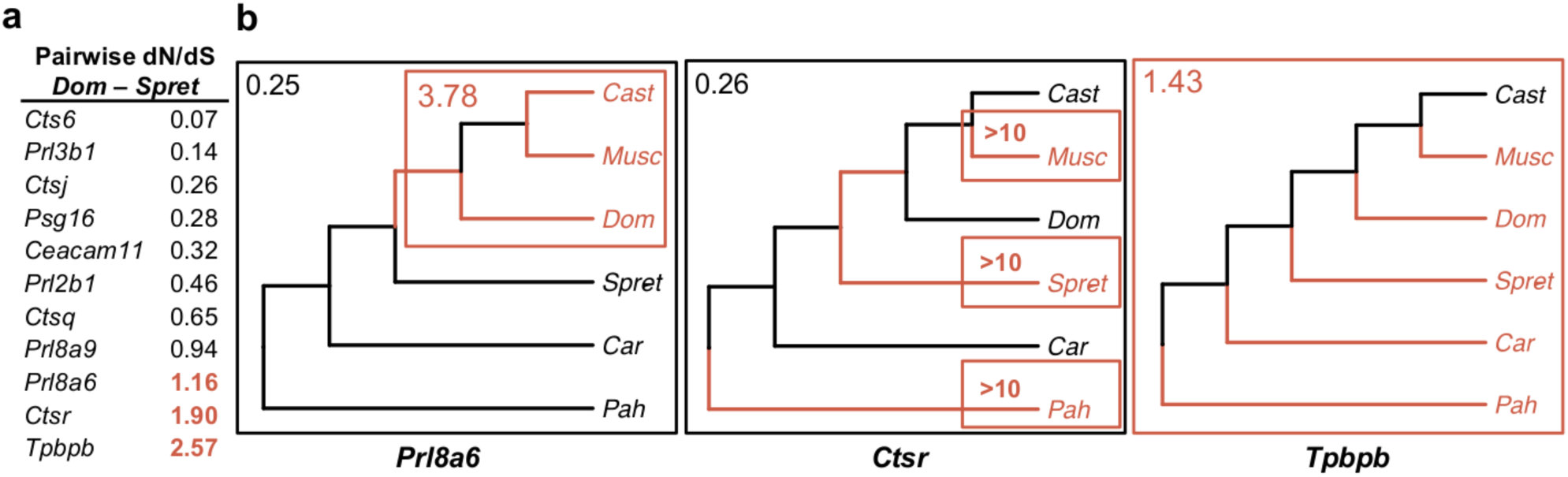
Evolutionary rates of placenta-specific gene family (PSF) genes expressed in the maternal medial preoptic area (MPoA). (A) Pairwise evolutionary rate between *Mus m. domesticus (Dom)* and *Mus spretus (Spret)* for *Ctsr* and the 10 co-expressed PSF genes with the highest expression in MPoA. Evolutionary rate (dN/dS) > 1 is marked in red throughout the figure and is indicative of positive selection. (B) PAML4 CodeML analysis results for the three genes with pairwise dN/dS>1. Species are *Mus m. castaneus (Cast), Mus m. musculus (Musc), Mus m. domesticus (Dom), Mus spretus (Spret), Mus caroli (Car), Mus pahari (Pah)*. dN/dS values are indicated for groups of branches depending on which CodeML model provided the best fit for the data (M0: one evolutionary rate for the whole tree, MC: selected branches evolve at a different rate than the rest of the tree). dN/dS values depicted in black for *Prl8a6* and *Ctsr* indicate background evolutionary rates.

For all three genes, we found evidence for positive selection on *Dom* (*Prl8a6*) or *Spret* branches (*Ctsr*), or both (*Tpbpb*) (Fig. 5, Supplemental Table S1). Specifically, there was evidence for positive selection on *Prl8a6* in the *Mus musculus* subspecies clade (LRT_(M0-MC)_=7.50, p=0.01, foreground dN/dS_(MC)_=3.78, background dN/dS_(MC)_=0.25). Evolutionary rates for *Ctsr* were elevated on branches leading to *Pah, Spret* and *Musc* relative to the rest of the tree (LRT_(M0-MC)_=6.93, p=0.01, foreground dN/dS_(MC)_>10, background dN/dS_(MC)_=0.26). Results for *Tpbpb* suggest high evolutionary rates across the whole tree (LRT_(M0-MC)_=3.56, p=0.1, dN/dS_(M0)_=1.43) (Fig. 5, Supplemental Table S1).

## Discussion

Molecular communication between the placenta and the maternal brain is crucial for the expression of maternal behavior in rodents (Bridges et al. 1996; Larsen and Grattan 2012; Creeth et al. 2018). In humans, altered regulation of placental imprinted genes is associated with prenatal depression (Janssen et al. 2016b), a predictor of lower growth rate and higher disease risk in infants (Rahman et al. 2004). In this study, we used a hybrid mouse model to characterize the extent to which placental disruption influences gene expression in the maternal brain. Several maternally expressed imprinted genes were transgressively misexpressed in the hybrid placenta. In *Mus m. domesticus* females carrying hybrid litters we found altered placenta-specific gene family expression in the placenta, and in the maternal MPoA. Surprisingly, the expression of these genes was highly correlated between the two tissues, and was *Mus spretus*-like in the MPoA. This suggests that paternally inherited alleles in the placenta exert substantial influence on expression in the maternal brain. Collectively, our results reveal reciprocal effects of mothers on offspring and offspring on mothers, mediated in both cases by the placenta. We discuss these findings in light of maternal-fetal coevolution and parental conflict, and identify potential implications for placental pathologies.

### Maternal effects on placental expression

Global patterns of expression in the placenta were strongly associated with maternal genotype. In hybrids, half as many genes were DE relative to normal *Dom* placentas as opposed to *Spret* placentas. Notably, the >1,600 genes with *Dom*-like expression in hybrid placenta were highly enriched for terms associated with immunity and regulation of blood flow, both of which are essential to placental mediation between mother and embryo (Cross et al. 1994). Because maternal vasculature is incorporated into the placenta, whole placenta transcriptomes necessarily include some transcripts of maternal origin and detection of *Tnfsrs23* transcripts in all samples indicates that some cells from the maternal decidua remained following dissection. However, maternal blood flow within the placenta is under the direct control of placental cell lineages; trophoblast giant cells invade and replace maternal vascular endothelium, limiting maternally-derived tissue to blood (Rai and Cross 2014). Likewise, *Tnfsrs23*-expressing cells are adjacent to fetal trophoblast giant cells at the junction between fetal and maternal tissues (Clark et al. 2002). Therefore, while contamination from maternal transcripts may contribute to *Dom*-like expression in hybrid placenta, it is unlikely to bias the expression of such a large number of genes. The regulatory effects of maternal hormones, and of maternally inherited genes in the placenta, are non-mutually exclusive alternative explanations. For example, because paternal X chromosome inactivation is maintained in mouse placenta (Tagaki and Sasaki 1975), maternally inherited X-linked genes are strong candidates for modulating autosomal expression in both sexes. While disentangling maternal effects (*sensu* Wolf and Wade 2009) from the effects of maternally inherited genes is a challenge for future studies, we note that the match between maternal genotype and placental expression of genes that modulate maternal immune tolerance and angiogenesis is consistent with the expectation of molecular coadaptation between mother and offspring (Wolf and Brodie 1998, Keverne and Curley 2008), and the well-established effect of maternal environment on placental function (Cottrell and Seckl 2009; Monk et al. 2012).

### Altered PSF and IG expression in the placenta

Maternal adaptation to pregnancy relies to a great extent on placental signaling. Thus, altered expression of genes encoding or influencing placental signaling molecules can ultimately affect maternal physiological and behavioral response to pregnancy (Bridges et al. 1996; Creeth et al. 2018). Misexpression in the hybrid placenta was substantial. However, the most striking pattern we found was the reduced expression of a large number of PSFs. In mice, most of these genes are expressed from the placental endocrine compartment and many are found in maternal plasma during pregnancy (Rawn and Cross 2008). In this hybrid mouse model, the endocrine compartment is markedly reduced when the mother is *Dom* (Zechner et al. 1996). Thus, reduced abundance of PSF producing cell types likely contributes to overall reduction in PSF expression.

IGs are thought to modulate PSF expression, primarily through effects on placental endocrine cell abundance, with maternally expressed genes (MEGs) repressing and paternally expressed genes (PEGs) promoting cell proliferation (John 2013, 2017). Two such MEGs, *Phlda2* and *Ascl2*, were transgressively misexpressed in hybrid placentas. Altered expression of either of these genes in lab mouse models results in an undersized endocrine compartment, altered glycogen energy stores and reduced PSF gene expression (Tunster et al. 2010, 2016a,b). Indeed, *Phlda2* and *Ascl2* seem to be critical co-regulators of placental endocrine compartment development (John 2017). Two other MEGs, *Dcn* and *Grb10*, were overexpressed when compared to *Dom* placentas. While neither is specifically implicated in placental endocrine function, both are key modulators of placental growth, and *Dcn* overexpression represses cellular proliferation (Yamaguchi and Ruoslahti 1988; Kresse and Schönherr 2001; Garfield et al. 2011). Collectively, our results are consistent with the proposed role of IGs in placental signaling (Haig 1996; Tunster et al. 2013; John 2013,2017), and identify MEG misexpression as a candidate mechanism for the undersized endocrine compartment and consequent global reduction in PSF expression in hybrid placenta.

### The effects of hybrid placental dysfunction on the maternal brain

Altered signaling in hybrid placentas has the potential to affect the maternal brain. We found subtle but significant differences in the expression of four genes in the MPoA of *Dom* females exposed to hybrid relative to conspecific placentas. Both *Fmn1* (Formin1) and *Caln1* (Calneuron1) were underexpressed. In the brain, *Fmn1* is involved in the formation of adherens junctions and in linear actin cable polymerization (Kobielak et al. 2004). The formation of adherens junctions is important in the maintenance of the blood brain barrier (BBB), a highly specialized structure that regulates the influx of molecules into the brain (Stamatovic et al. 2008). During pregnancy, the permeability of the BBB is increased by placenta-derived factors to which the maternal brain must respond in order to maintain this barrier (Cipolla 2007; Schreurs et al. 2012). Reduced expression of *Fmn1* therefore suggests alterations in BBB adaptation during hybrid pregnancies. *Caln1* encodes a neuron-specific protein with sequence similarities to calcium-binding calmodulins. While the function of *Caln1* is uncharacterized, homology to calmodulin suggests a role in neuronal calcium signaling (Wu et al. 2001). Decreased expression of a calcium-binding protein could indicate alterations in neuronal activity in the MPoA exposed to hybrid placentas.

*Drd3* (Dopamine receptor D3) was also underexpressed compared to *Dom* mothers, but not to *Spret* mothers, in the hybrid pregnancy MPoA. DRD3, a D2-like receptor with a generally inhibitory role, is implicated in treatment-resistant major depression (Lattanzi et al. 2002) and *Drd3* knock-out mice exhibit a suite of anxiety- and depressive-like behaviors with similar but milder effects in heterozygous knock-outs (Accili et al. 1996; Moraga-Amaro et al. 2014). Given that the action of dopamine in the MPoA is critical for the expression of maternal behavior in rats (Numan and Stolzenberg 2009), and hypothalamic dopamine is altered in a mouse model for post-partum depression (Avraham et al. 2017), reduced *Drd3* expression in the MPoA might cause deficits in maternal behavior. However, *M. spretus* mothers are more responsive to pups than *M. m. domesticus* mothers (Cassaing et al. 2010) and *Drd3* expression in the hybrid pregnancy was statistically indistinguishable from the normal *Spret* pregnancy. How, and to what extent placental expression of paternally inherited alleles promotes maternal behaviors is an intriguing question for in-depth studies.

*Ctsr* (Cathepsin R), a placenta-specific cathepsin, was the only gene that was overexpressed in the MPoA exposed to hybrid placentas. The difference in expression, although significant, was so small that the biological relevance is questionable. However, unlike other PSF genes, expression of *Ctsr* in the maternal brain was unique to the hybrid pregnancy. Interestingly, loss of the IG *Peg3* leads to de-repression of several PSF members, including *Ctsr*, in the fetal and adult brain (Kim et al. 2013). While *Peg3* was not misexpressed in the hybrid placenta, the transgressively overexpressed MEG, *Phlda2*, was recently shown to perturb maternal behavior and neural gene expression when its dosage was altered in mouse placenta (Creeth et al. 2018). Specifically, overexpression of placental *Phlda2* reduced postpartum nurturing behavior, while underexpression increased maternal behavior (Creeth et al. 2018). Since *Phlda2* and *Ascl2* jointly regulate development of the endocrine compartment (John 2017), it is likely that transgressive misexpression of both genes in hybrid placenta impacts the maternal brain via effects on placental hormone expression.

### Paternal effects on the maternal brain

The hybrid placenta expresses both maternally derived (*Dom*) and paternally derived (*Spret*) alleles. Thus, females pregnant with hybrids are exposed to gene products from a foreign paternal genome. In *Dom* females exposed to hybrid placentas we found a substantial subset of genes, including PSF genes and *Drd3*, with expression patterns that differed from *Dom* mothers with conspecific litters, but closely matched those of *Spret* mothers.

A surprisingly large number of genes were co-expressed between placenta and MPoA in hybrid and *Spret* pregnancies but not in *Dom* pregnancies. In particular, placental and MPoA PSF gene expression was highly correlated in hybrid pregnancies and in *Spret* pregnancies, while *Dom* pregnancies showed a weaker correlation. Likewise, total MPoA PSF gene expression was *Spret*-like in hybrid pregnancies. These results suggest that PSF expression levels in the maternal MPoA are driven by placental expression levels of the same genes, and that PSF, and potentially *Drd3*, expression in the MPoA is influenced by paternally inherited alleles in the placenta. Thus, natural differences between the parental species used in this study reveal an unanticipated effect of the paternal genome on the maternal brain. This hypothesis is illustrated in Fig. 6.

**Figure 6.**
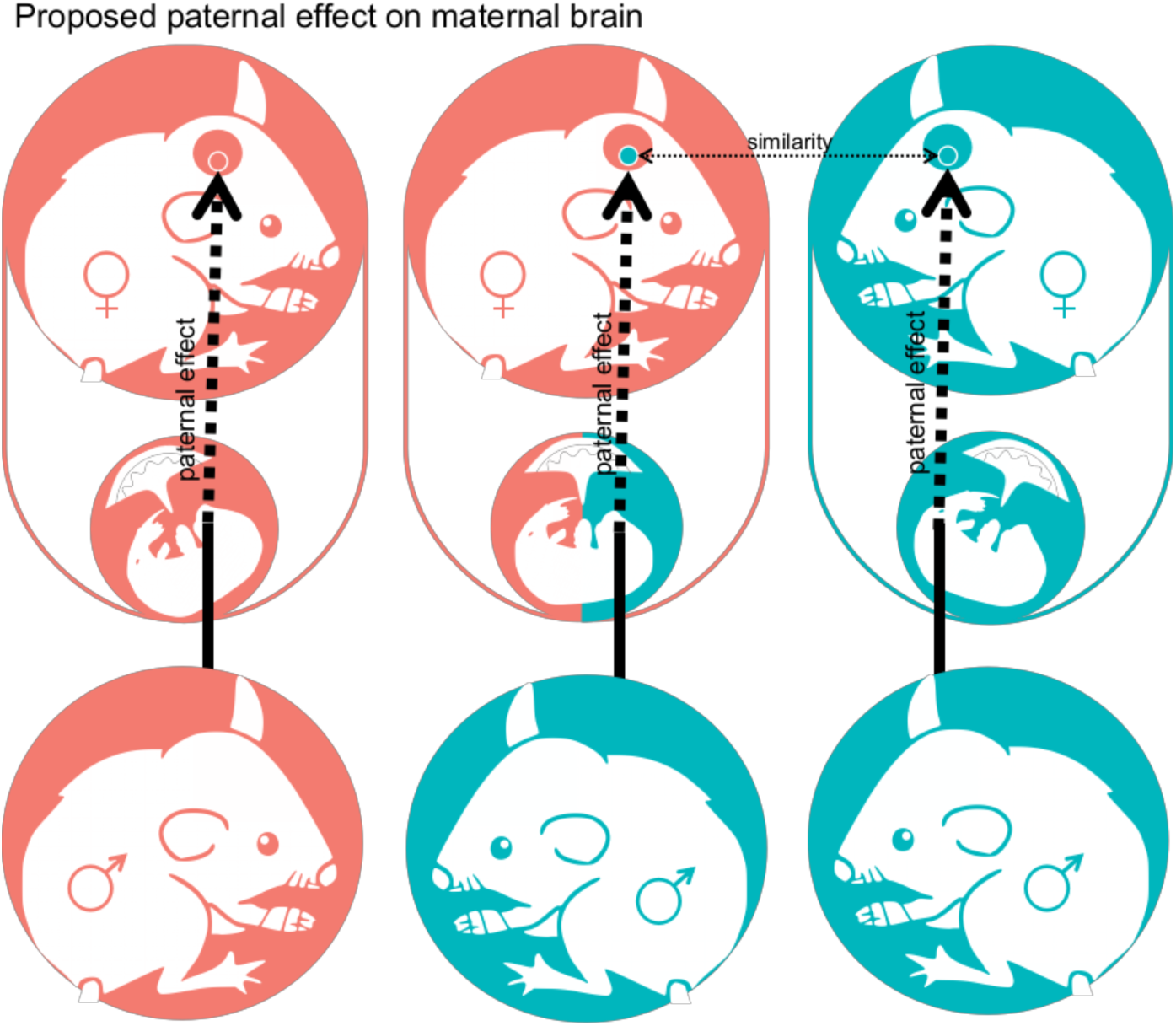
Proposed paternal effect on maternal brain. Schematic visualization of the three different crosses and the potential indirect effect of the paternally inherited allele through the placenta on the maternal brain. *Mus m. domesticus* is depicted in red and *Mus spretus* in blue. Hybrid tissue is indicated by a combination of red and blue.

A reciprocal shift towards *Dom*-like PSF expression in the brains of *Spret* females carrying litters sired by *Dom* males would provide strong support for this hypothesis. Unfortunately, generating hybrid pregnancies in this direction of the cross is notoriously difficult (Zechner et al. 1996) and our attempts failed (see Methods). However, the generality of paternally-mediated placental effects on mothers could be tested in other species crosses characterized by parent-of-origin-dependent placental abnormalities (e.g. *Phodopus* hamsters (Brekke and Good 2014) and *Peromyscus* mice (Vrana et al. 1998)).

### A signature of conflict in PSF evolution and expression in the maternal brain

Pregnancy requires substantial investment from the mother, which is offset by costs to her capacity to invest in future offspring (Trivers 1972). However, when offspring are sired by multiple males, selection favors fathers who extract maximal maternal resources for their own offspring (Trivers 1972). Haig and colleagues proposed that these asymmetries in the reproductive interests of males and females, and the coefficients of relatedness between mothers and offspring (always 0.5) vs. fathers and offspring (0.5 or 0), should promote parental antagonism, played out at the molecular level between maternally and paternally expressed IGs in the placenta (Moore and Haig 1991; Haig and Graham 1991; Haig 2000). Because placental endocrine signals promote maternal investment in current offspring, placental hormones are also proposed players in both parental and mother-offspring conflicts (Trivers 1985; Haig 1996). Consistent with a history of antagonistic coevolution, PSFs in general are the fastest evolving genes in the rodent placenta (Chuong et al. 2010). We report a similar signature of selection on three PSF genes that are co-expressed in the hybrid placenta and the maternal MPoA.

Trivers (1985) described placental hormones as the molecular equivalent of begging calls. Here we show for the first time that the expression of Prls and other PSFs is highly correlated between placenta and maternal brain. While the function of PSFs in the brain is undefined, placental genotype-dependent differences between *Dom* females in the strength of the correlation and the number of co-expressed genes indicate that the relationship is driven by the placenta not the mother. Moreover, the *Spret*-like co-expression patterns of PSF genes in mothers of litters sired by *Spret* males implicate the paternally inherited genome as the driver of these placental begging calls, which are echoed in the maternal brain. Given that these patterns of expression are consistent with parental conflict, it is noteworthy that the opportunity for sperm competition (inferred from testis-body mass ratios) is higher in *M. spretus* than in *M. m. domesticus* (Gomez Montoto et al. 2011).

### Preeclampsia related gene expression in hybrid pregnancies

Preeclampsia is a serious pregnancy complication and the lead cause of maternal and fetal morbidity and mortality (Burton and Jauniaux 2004; Redman and Sargent 2009). We found significant overlap between transgressively misexpressed genes in hybrid placentas and preeclampsia related genes and pathways. Haig (1993) interpreted preeclampsia as a consequence of conflicts between maternal and paternal genomes, played out in the placenta and potentially involving IGs. While the genetic basis of preeclampsia is complex (Uzun et al. 2016), involvement of IGs is supported in both humans and mouse models. For example, human chromosome 10 regions containing imprinting clusters are associated with preeclampsia (Oudejans et al. 2004), and loss of the MEG *Cdkn1c* causes preeclampsia-like symptoms in mice (Kanayama et al. 2002). Interestingly, *Cdkn1c* is in the same imprinting cluster as *Phlda2* and *Ascl2* (dist7, IC2), both of which were misexpressed in hybrid placentas. While *Cdkn1c* was not differentially expressed in the hybrid placenta, it is possible that misexpression in this imprinting cluster is a general contributor to preeclampsia-like placental phenotypes. Because preeclampsia can significantly alter permeability of the BBB (Cipolla 2007), it is also notable that *Fmn1*, a gene implicated in BBB maintenance, was underexpressed in brains of mothers exposed to hybrid placentas.

## Conclusions

Evolutionary theoreticians have modeled mammalian pregnancy as both intimate cooperation and antagonistic struggle between two genetically distinct organisms (Trivers 1974; Haig 1993; Wolf and Hager 2006). Whether driven by conflict or coadaptation, it is clear that the placenta is the mediator of these complex interactions between mother and offspring. Here we concentrated on placental effects on the maternal brain during the final stages of pregnancy, when it is believed to be a critical source of signal molecules that prime female physiology and behavior for motherhood (Larsen and Grattan 2012; Bridges 2015). We found both hybrid placental misexpression with the potential to disrupt maternal-fetal communication, and altered expression in the brains of mothers exposed to hybrid placentas. Expression in the hybrid placenta seems to be dominated by the maternally derived genome and/or driven by maternal effects. Maternal-placental communication genes co-expressed in maternal brain and placenta show elevated evolutionary rates, consistent with antagonistic coevolutionary processes. The expression of a proportion of transcripts of these genes from a foreign paternal genome in the placenta has the potential to affect the maternal brain and alter maternal behavior. In addition to the effects of placental disruption on the maternal brain, natural differences between the parental species in this hybrid system suggest a previously undescribed influence of the paternal placental genome on the maternal brain. These paternal effects on the maternal brain could play a major role in the expression of maternal behavior and the quality of maternal care, and open novel avenues of research in both evolutionary and biomedical fields.

## Methods

### Animals and tissue collection

Mice used in this study were maintained on a 12:12 light:dark cycle with lights on at 9:00 AM, and were provided with 5001 Rodent Diet (LabDiet, Brentwood, MO, U.S.A.) and water ad lib. All animal procedures were approved by the Oklahoma State University IACUC under protocol #141-AS. *Mus m. domesticus* (*Dom*) was represented by the wild-derived inbred strain WSB/EiJ (Jackson Laboratory) and *Mus spretus* (*Spret*) was represented by the wild-derived inbred strain SFM/Pas (Montpellier Wild Mice Genetic Repository). We conducted three crosses (female shown first): *Dom* X *Dom* (n=5), *Dom* X *Spret* (n=5), *Spret* X *Spret* (n=5). The reciprocal cross (*Spret* X *Dom*) was attempted 25 times but was never successful. Prior to pairing, females were placed in a cage with soiled conspecific male bedding for ∼48 hrs to induce receptivity to mating (Whitten 1956). Mice were paired between 5:00 and 6:00 PM, left undisturbed for two nights, and split on the morning of the second day. The second night was counted as embryonic day 0 (e0). Females were weighed after two weeks to confirm pregnancy but were otherwise left undisturbed. Pregnant females (n=5/type of pregnancy) were euthanized by cervical dislocation between 10:00 and 11:00 AM on embryonic day 17-18 (e17.5) and the maternal brain was extracted. Embryos were separated from placentas and placentas were weighed, and the maternally-derived decidual layer was removed as previously described (Qu et al. 2014) and discarded. Only fetally-derived placental tissue was used in this study. All tissues were transferred immediately to RNAlater (Thermo Fisher, USA), kept at 4°C overnight to allow RNAlater perfusion, and stored at −20°C until microdissection and RNA extraction.

### Brain microdissection and RNA extraction

The maternal MPoA was localized using the Mouse Brain atlas (Figs. 26-33, (Paxinos and Franklin 2013)), and microdissected by sectioning the RNAlater perfused brain at 100µm on a Leica CM 1950 cryostat, followed by dissection under a dissecting microscope in chilled PBS droplets for improved visibility of brain microstructure. DNA was extracted from embryonic tissue using the DNeasy Blood & Tissue Kit (Qiagen, USA) followed by PCR for the Y-linked gene, *Zfy1*, to determine sex. Placentas from one male and one female per litter were used for RNA extraction (n=5 males/cross, n=4 females/hybrid cross, n=5 females/conspecific cross). RNA was extracted from all tissues immediately after microdissection using the RNeasy Plus Universal Mini Kit (Qiagen) for MPoA, and the AllPrep RNA/DNA Mini Kit (Qiagen) for placenta. RNA was stored at −80°C until sequencing.

### RNAseq pipeline

#### Sequencing (RNAseq)

RNA integrity (RIN) was determined by the sequencing facility (Novogene, Sacramento, CA) using the RNA Nano 6000 Assay Kit with the Agilent Bioanalyzer 2100 system (Agilent Technologies, Santa Clara, CA, USA). RIN values ranged from 8.2-10 for all samples. Library preparation was performed by the sequencing facility, using the NEB Next Ultra RNA library prep kit for Illumina. RNAseq was performed on the Illumina HiSeq 4000 platform, producing >30 million, 150bp paired-end reads per sample.

#### Mapping

QC of raw sequencing reads and trimming were performed in Trim Galore! 0.4.5 (Brabraham Bioinformatics, http://www.bioinformatics.babraham.ac.uk/projects/trim_galore), using a phred score cutoff of 30 and minimum sequence length of 100 after trimming. In order to map hybrid placenta reads we generated a pseudo-hybrid genome using the genome preparation tool of the program SNPsplit (Brabraham Bioinformatics, (Krueger and Andrews 2016)). Briefly, SNPs from both *Dom* (WSB/EiJ) and *Spret* (SPRET/EiJ) relative to the mouse genome (GRCm38.89) available from the Ensembl FTP server (ftp://ftp.ensembl.org) were introduced into the mouse genome (GRCm38.89). SNPs between *Dom* and *Spret* were then N-masked to allow mapping of both *Dom*- and *Spret*-derived reads. To improve comparability, all placenta samples (*Dom*, *Spret* and hybrid) were mapped to the pseudo-hybrid genome. MPoA samples were mapped to their corresponding genomes (WSB/EiJ_v1 for *Dom* MPoA, SPRET/EiJ_v1 for *Spret* MPoA, (Keane et al. 2011)). Mapping was done using HISAT2 2.1 (Kim et al. 2015). After mapping we filtered the resulting alignment files using SAMtools 0.1.19 (Li et al. 2009), retaining only high quality (HISAT2 MAPQ score 40), uniquely mapped, paired reads for analysis.

#### Post-processing of alignments

Before filtering, average alignment rate for MPoA samples was 87% for *Spret* MPoA samples and 88% for *Dom* MPoA samples. For placenta samples the alignment rate was slightly lower for *Spret* samples (88%) compared to *Dom* (91%) and *hybrid* (91%). We therefore randomly downsampled all alignment files to ∼40 million reads using SAMtools 0.1.19 (Li et al. 2009) to account for a possible mapping bias and to improve comparability.

Allele-splitting of hybrid alignments: To confirm ∼equal expression from *Dom* and *Spret* alleles in the hybrid placenta we split the hybrid alignment files, separating reads originating from the *Spret* and *Dom* alleles. Splitting was performed using the program SNPsplit (Brabraham Bioinformatics, (Krueger and Andrews 2016)). The pseudo-hybrid genome was produced by introducing SNPs between both Spret and mouse and Dom and mouse into the mouse genome and then N-masking those SNPs that are present between Dom and Spret. Mapped reads can therefore be assigned to the *Dom* or *Spret* allele based on the list of SNPs between *Dom* and *Spret* and overlap with N-masked positions in the mouse genome. Using this procedure, SNPsplit produced separate alignment files for each allele containing clearly assignable reads. These were then quantified and used to produce a diagnostic boxplot, demonstrating ∼equal expression of both alleles in the hybrid placenta (Supplemental Fig. S8).

#### Quantification

Transcript quantification and annotation was done using StringTie 1.3.3 (Pertea et al. 2015). Gene annotation information was retrieved from the Ensembl FTP server (ftp://ftp.ensembl.org) for *Spret* (SPRET/EiJ_v1.86) and *Dom* (WSB/EiJ_v1.86). Mouse genome annotation was used for samples mapped to the pseudo-hybrid genome (GRCm38.89). We used the python script (preDE.py) included in the StringTie package to prepare gene-level count matrices for analysis of differential gene expression.

#### Differential expression (DE) analysis

Differential expression was tested with DESeq2 1.16.1 (Love et al. 2014). Pseudogenes were removed from the count matrices based on “biotype” annotation information extracted from Biomart (R-package biomaRt, (Durinck et al. 2005)). Low counts were removed by the independent filtering process implemented in DESeq2 (Bourgon et al. 2010). The adjusted p-value (Benjamini-Hochberg method) cutoff for DE was set at 0.05. Due to variation in litter size, especially in females carrying hybrid litters (range=3-6), and its potential effect of on MPoA expression, we corrected for litter size in all MPoA sample comparisons. To analyze DE between *Spret* and *Dom* MPoA, which were mapped to their respective genomes, we extracted homologous gene names from the mouse genome database using Ensembl Biomart (R-package biomaRt, (Durinck et al. 2005)) and merged the dataset based on the genes that had a clear mouse homolog in both. Normalized read count tables produced by DESeq2 were used in subsequent co-expression analyses.

#### Co-expression

To determine co-expression between placenta and MPoA we set a cutoff of 10 normalized counts for at least 4 out of 5 observations each tissue type (MPoA, male placenta and female placenta). Based on this cutoff we report genes expressed in both placenta and MPoA for each type of pregnancy. We then determined differences in co-expression between the three pregnancy types.

#### Gene ontology (GO) term and pathway overrepresentation analysis

We performed GO term and pathway overrepresentation analyses on relevant lists of genes from DE and co-expression analyses using the PANTHER gene list analysis tool with Fisher’s exact test and FDR correction (Mi et al. 2017). We tested for overrepresentation based on the GO annotation database (Biological Processes) (released 07-Jan-2017, (Ashburner et al. 2000; The Gene Ontology Consortium 2017)) and the Reactome pathway database (version 58 (Fabregat et al. 2017)).

### Evolutionary rates for selected genes

We extracted the pairwise evolutionary rate (dN/dS = nonsynonymous to synonymous substitution rate ratio) between *Dom* and *Spret* from Biomart (R-package biomaRt, (Durinck et al. 2005)). dN/dS is an index of selective pressure on coding sequence, with dN/dS = 1 indicating neutral evolution, dN/dS < 1 purifying selection, and dN/dS > 1 diversifying positive selection (Goldman and Yang 1994). Further analysis of genes with dN/dS > 1 (*Prl8a6*, *Ctsr*, *Tpbpb*) was performed with CodeML implemented in PAML 4.8 (Yang and Rannala 1997; Yang 2007), including sequences from related *Mus* subspecies and species (*Mus m. musculus (Musc)*, *Mus m. castaneus (Cast)*, *Mus caroli (Car)* and *Mus pahari (Pah)*). Coding sequences for all three genes for *Musc*, *Cast* and *Car* were available from Ensembl. For *Pah*, coding sequences for *Prl8a6* and *Ctsr* were downloaded from NCBI Genbank. For *Tpbpb*, we ran blastn on NCBI with the *Dom Tpbpb* coding sequence against the nr/nt database and found two matches for *Pah*, of which one showed higher similarity to *Dom Tpbpa* and the other to *Dom Tpbpb*. The latter was included in the CodeML analysis (Supplemental Table S5). CodeML calculates evolutionary rates by applying different models to an alignment and a phylogeny. To prepare the alignments, sequences were visualized with Geneious 9.1.8 (Biomatters, http://www.geneious.com/) and trimmed to coding sequence. Translation alignments were performed using the MUSCLE alignment algorithm, implemented in Geneious. The phylogeny was built based on recent phylogenomic analyses of house mice and related species (White et al. 2009; Sarver et al. 2017). For the CodeML codon frequency setting we used the setting with the best fit for each analysis according to the preliminary likelihood ratio analysis.

To calculate individual evolutionary rates for each branch in the tree we used CodeML’s “free-ratio” model. This served as an initial indication as to which branches might show higher evolutionary rates. After this, two models were computed: Model M0 “one ratio” in which all branches were constrained to evolve at the same rate and MC “two-ratio” in which selected branches are allowed to evolve at a different rate than the rest of the tree. Branches with potentially higher evolutionary rate based on the “free-ratio” model result were marked as foreground branches and were allowed to evolve differently from the background. To test if MC provides a better fit for the data than M0 we performed Likelihood Ratio Tests. When MC provided the better fit, and dN/dS calculated for the foreground branches was > 1 and dN/dS calculated for the background branches was < 1, we inferred positive selection on the foreground branches. When M0 provided the better fit and dN/dS for the whole tree was > 1, we inferred positive selection for the whole tree (Yang 1998).

### Data Access

RNA-seq data from this study have been submitted to the NCBI Gene Expression Omnibus (GEO; https://www.ncbi.nlm.nih.gov/geo/) under accession number GSE126469.

## Acknowledgements

We thank members of the Campbell lab for useful discussion, and B. Horn and S. Windle for assistance with mouse husbandry, and M. Nightengale for assistance in the lab. P.C. thanks K. Mack for generously sharing the ideas that catalyzed this project and J. Good for the gift of the SFM/Pas strain. This work was supported by the National Science Foundation (NSF-IOS 1558109 to P.C).

