## Supplemental Material for "Placental effects on maternal brain revealed by disrupted placental gene expression in mouse hybrids"

**Supplemental material in this pdf:**

Supplemental Fig. S1: Diagnostic plots for DESeq2 analysis for hybrid vs. Dom placenta.
Supplemental Fig. S2: Diagnostic plots for DESeq2 analysis for hybrid vs. Spret placenta.
Supplemental Fig. S3: Diagnostic plots for DESeq2 analysis for Dom vs. Spret placenta.
Supplemental Fig. S4: PCA diagnostic plot for Dom, hybrid and Spret placenta.
Supplemental Fig. S5: Diagnostic plots for DESeq2 analysis for MPoA-hy vs. MPoA-dom.
Supplemental Fig. S6: Diagnostic plots for DESeq2 analysis for MPoA-hy vs. MPoA-spret.
Supplemental Fig. S7: Diagnostic plots for DESeq2 analysis for MPoA-dom vs. MPoA-spret.
Supplemental Fig. S8: Allelic expression in hybrid placenta
Supplemental Table S4: PAML4 CodeML likelihood ratio test results.
Supplemental Table S5: Sequence ID information for evolutionary analysis.

**Additional supplemental material in separate files:**

Supplemental Dataset S1: Result summary and gene lists for placental DESeq2 analysis.
Supplemental Dataset S2: Result summary and gene lists for MPoA DESeq2 analysis.
Supplemental Dataset S3: Result summary and gene lists for placenta-MPoA co-expression analysis.

Figure S1. Diagnostic plots for DESeq2 differential expression analysis for hybrid vs. *Dom* placenta.

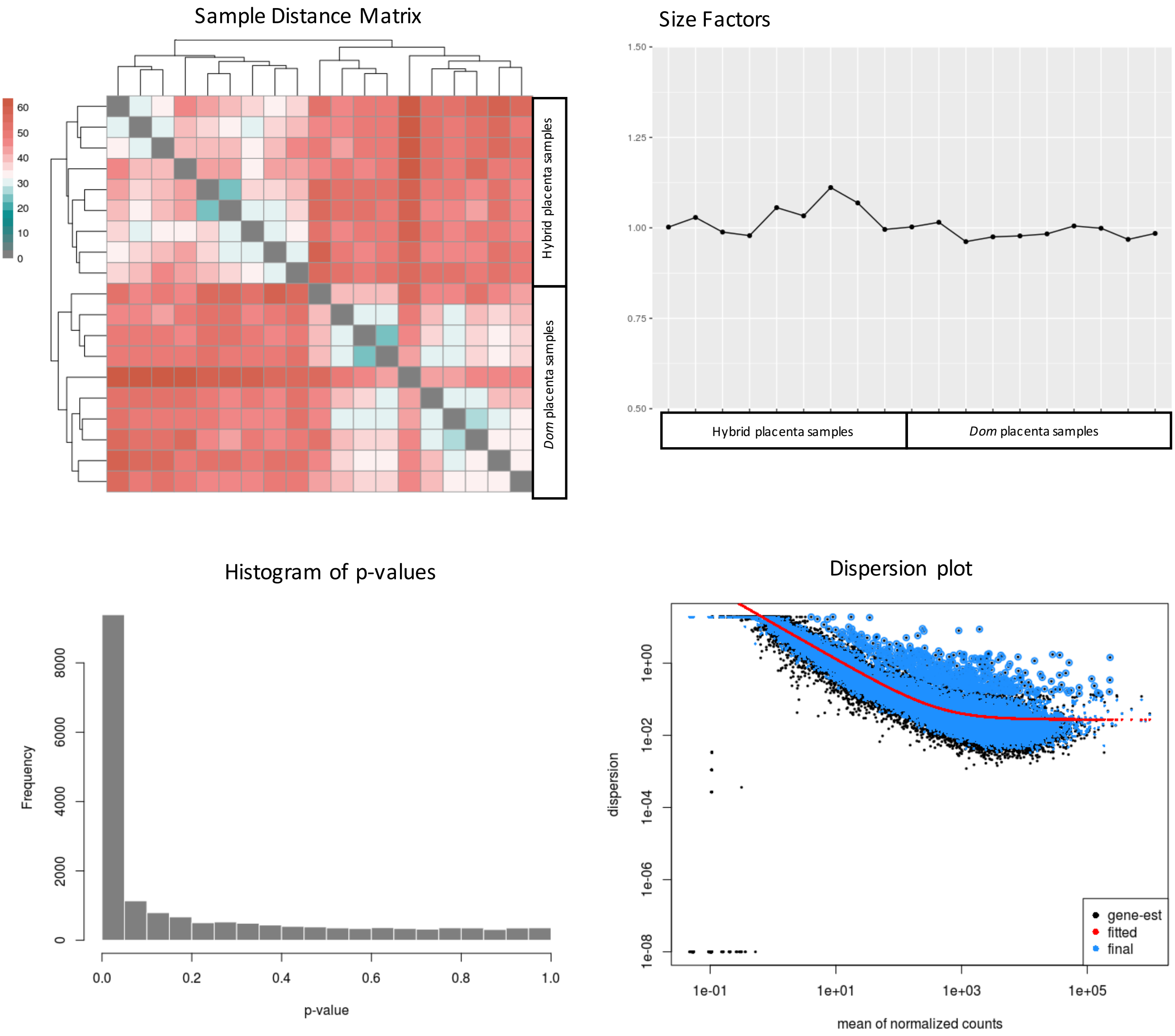

Figure S2. Diagnostic plots for DESeq2 differential expression analysis for hybrid vs. *Spret* placenta.

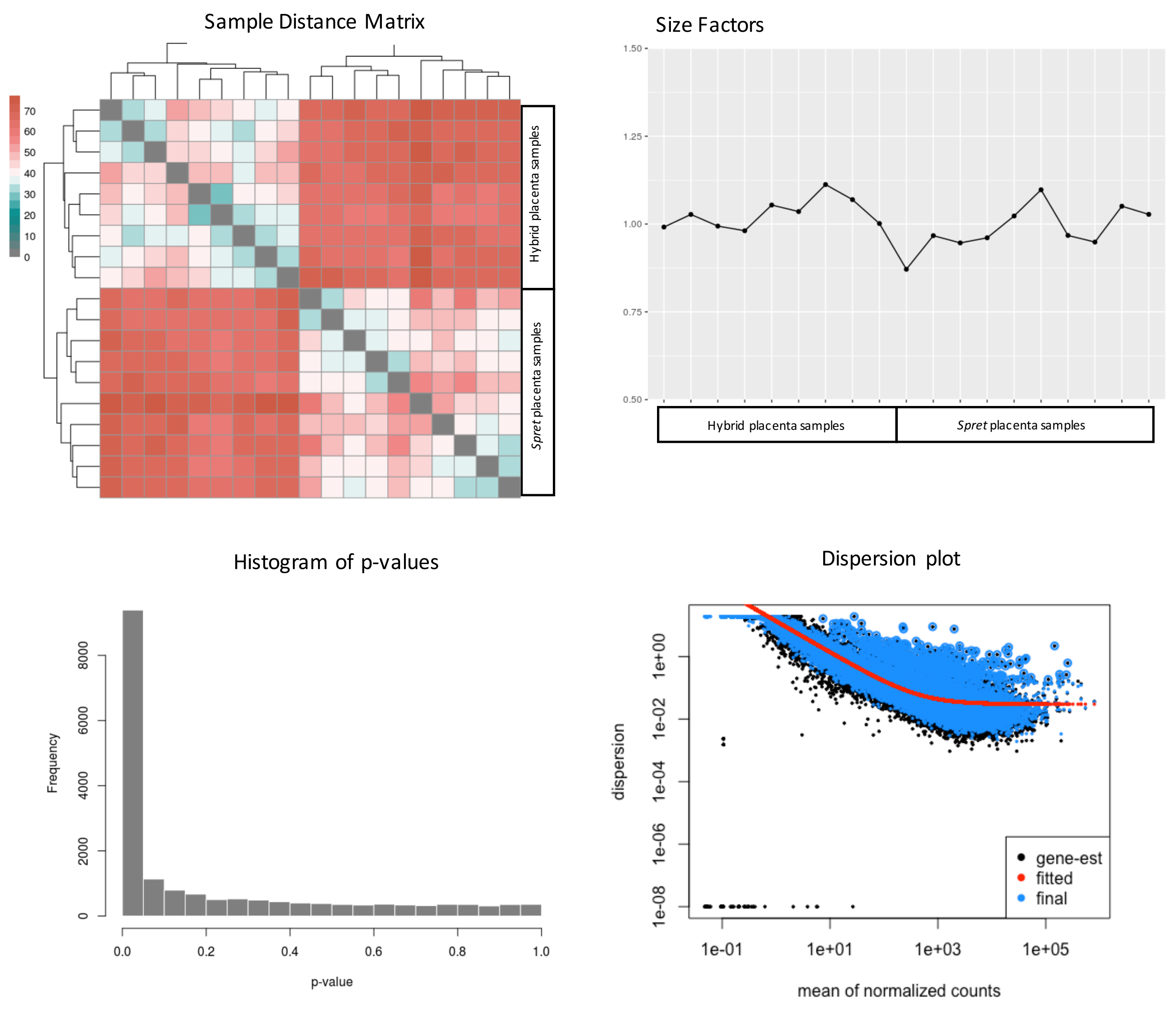

Figure S3. Diagnostic plots for DESeq2 differential expression analysis for *Dom* vs. *Spret* placenta.

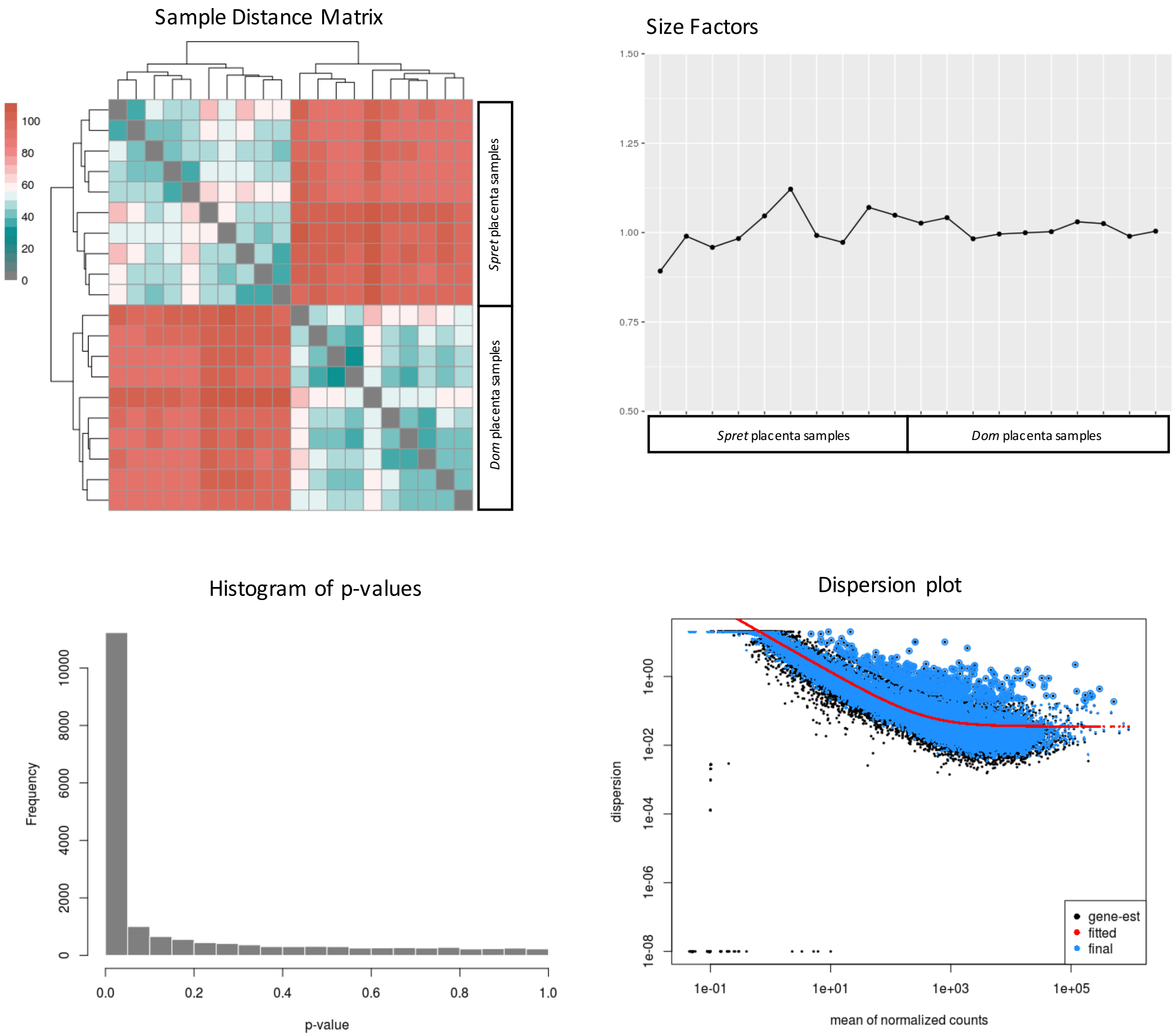

Figure S4. PCA diagnostic plot for *Dom*, *hybrid* and *Spret* placentas

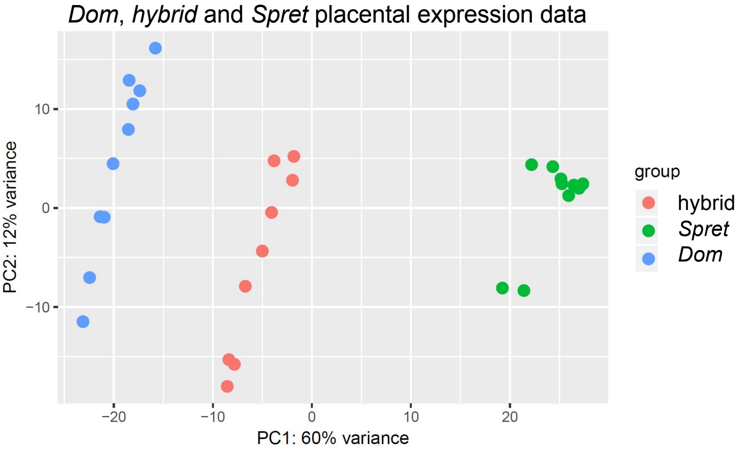

Figure S5. Diagnostic plots for DESeq2 differential expression analysis for MPoA-*hy* vs. MPoA-*dom*.

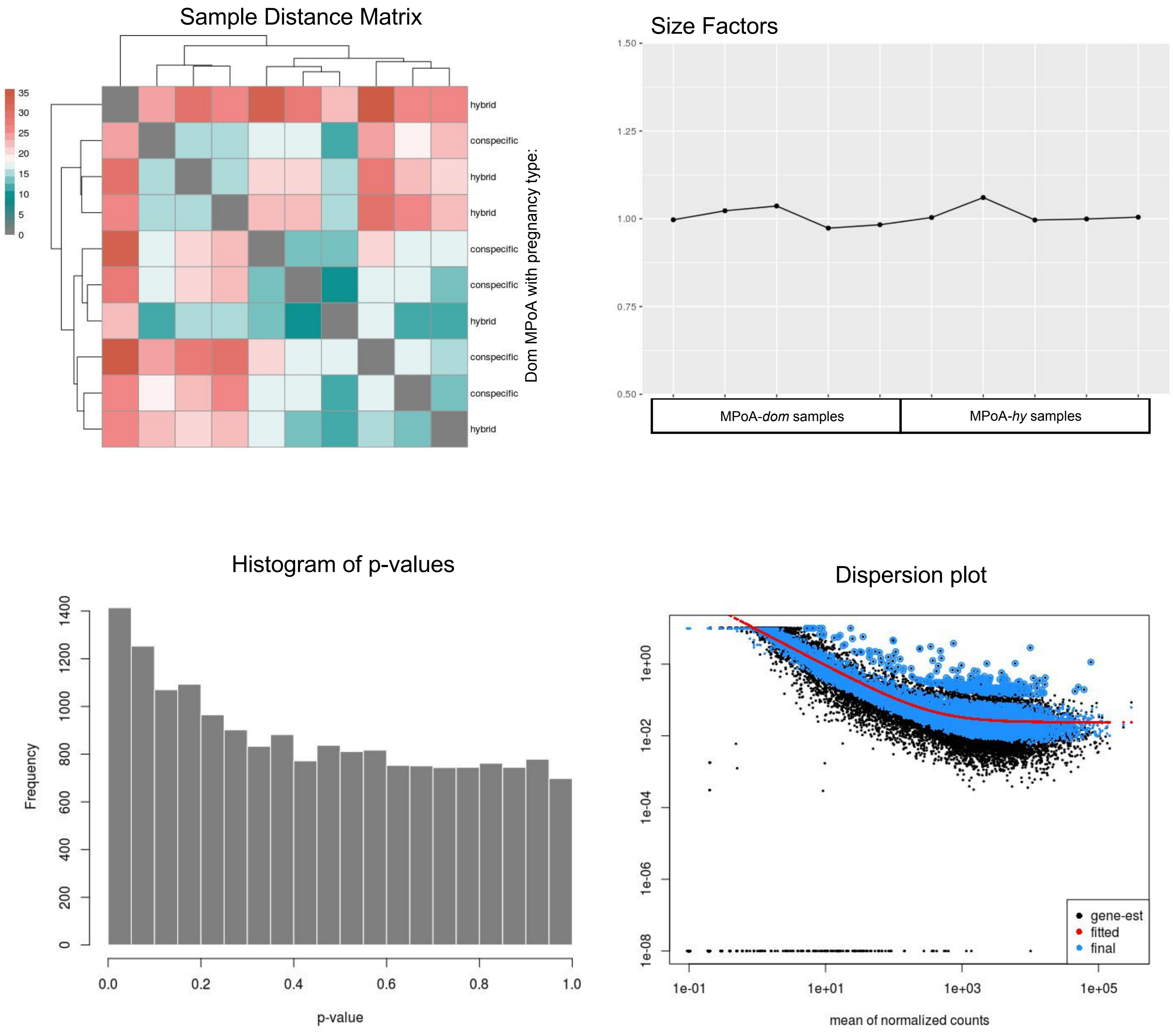

Figure S6. Diagnostic plots for DESeq2 differential expression analysis for MPoA-*hy* vs. MPoA-*spret*.

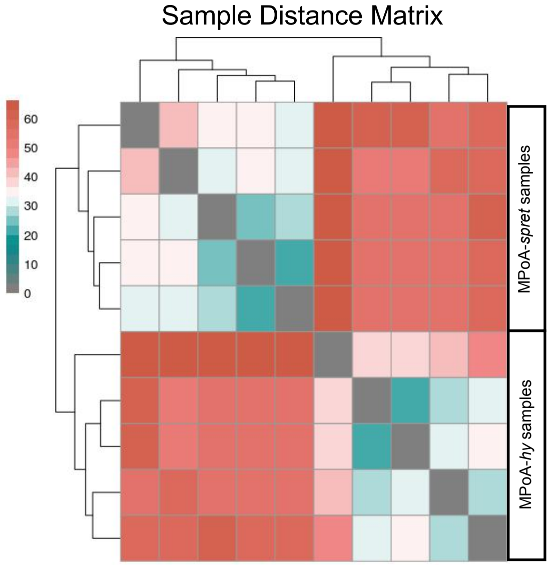


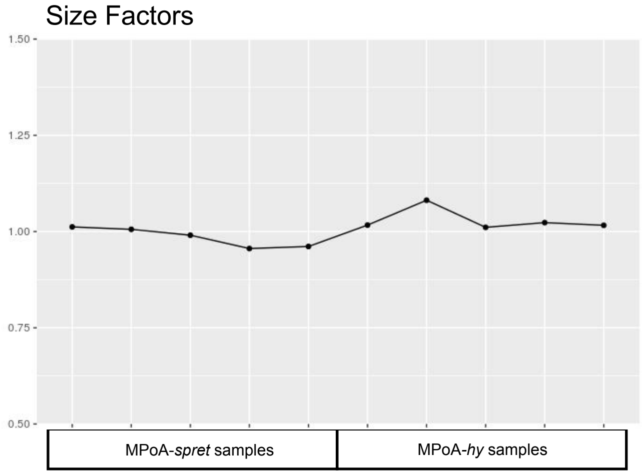


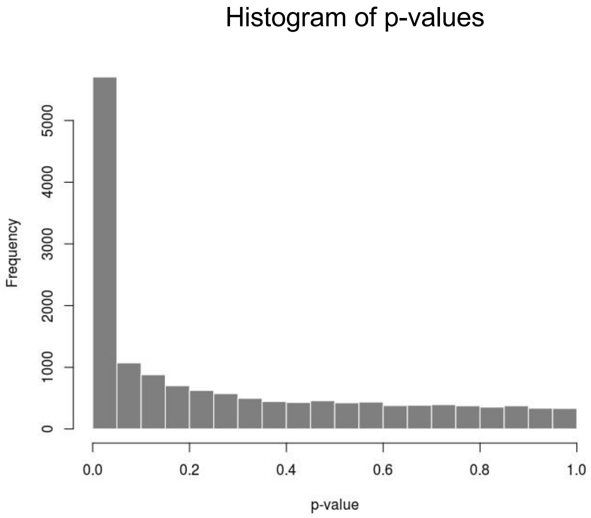


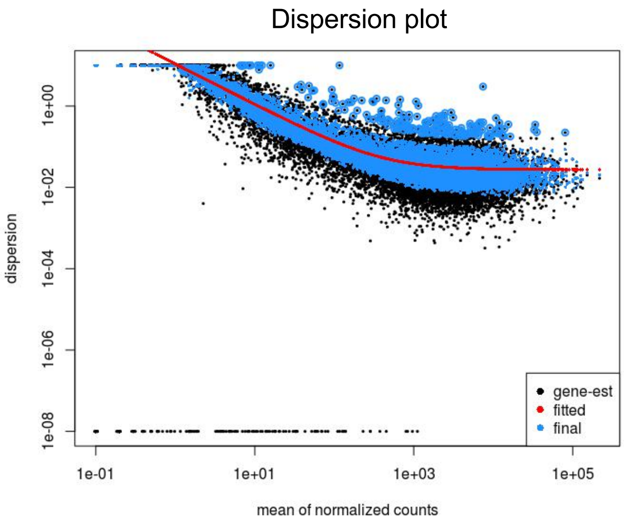

Figure S7. Diagnostic plots for DESeq2 differential expression analysis for MPoA-*dom* vs. MPoA-*spret*.

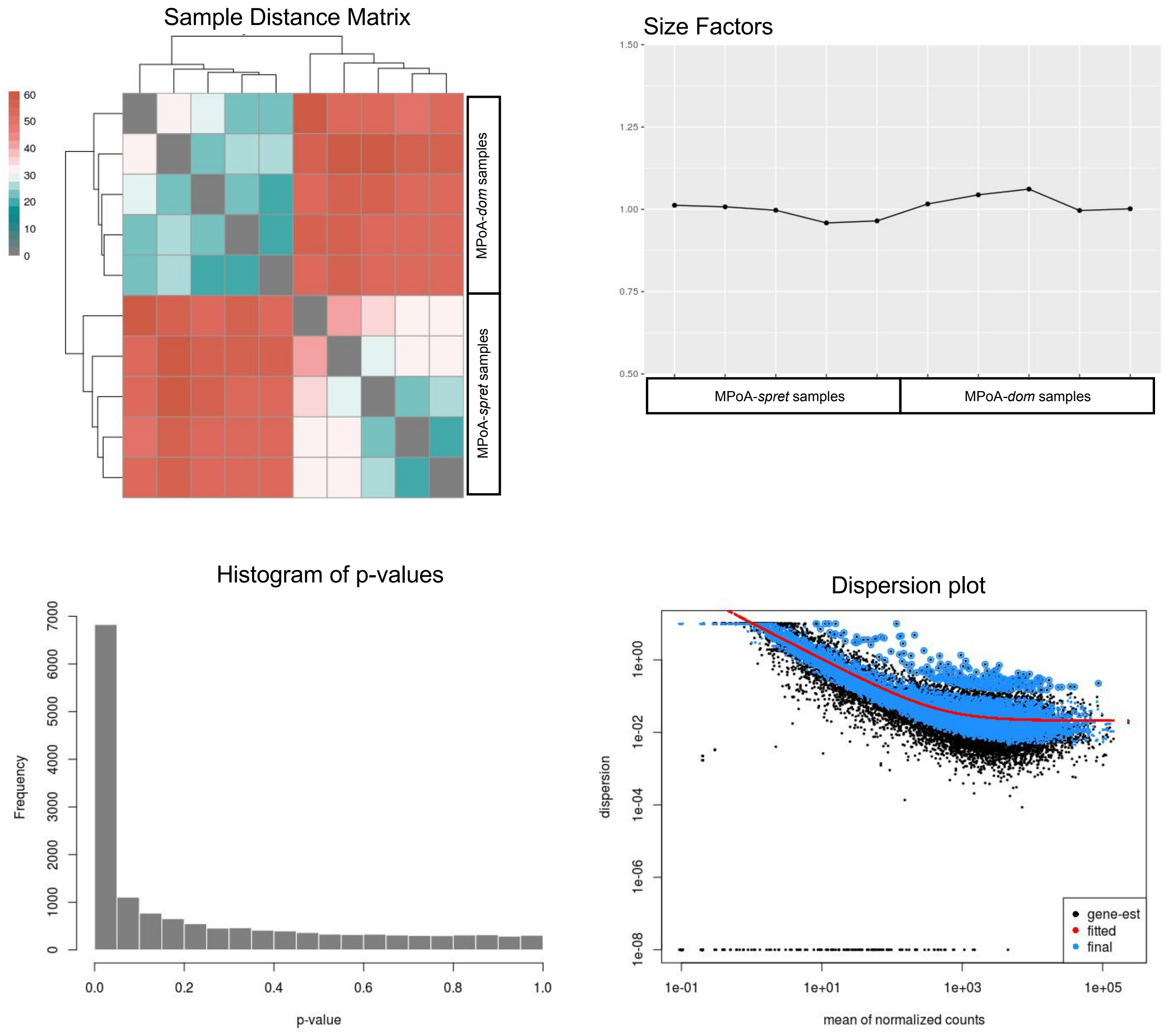

Figure S8. Allele expression in hybrid placentas

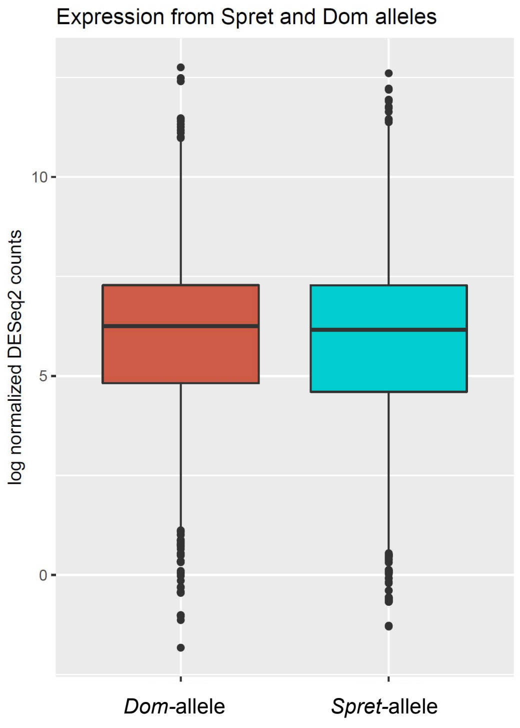

Table S1. PAML4 Codeml likelihood ratio test results

| | M0 lnL | MC lnL | 2Δ(M0-MC) | *p* | interpretation | dN/dS | Foreground dN/dS | Background dN/dS |
|---|---|---|---|---|---|---|---|---|
| *Ctsr* | 1380.921675 | 1377.454646 | 6.93405800000028 | 0.01 | differing rates across tree | M2 | >10 | 0.26 |
| *Prl8a6* | 1164.593902 | 1160.842685 | 7.50243399999999 | 0.01 | differing rates across tree | M2 | 3.78 | 0.25 |
| *Tpbpb* | 754.456437 | 752.675798 | 3.56127800000013 | 0.1 | whole tree evolves at same rate | M0 | 1.43 | |

M=Model, for explanation of compared models see methods, lnL=log likelihood of Model fit, dN/dS nonsynonymous to synonymous substitution rate ratio (evolutionary rate)

Table S2. Sequence ID information for evolutionary analysis

| | *Ctsr* | *Prl8a6* | *Tpbpb* | *Tpbpa* |
|---|---|---|---|---|
| *Musc* | MGP_PWKPhJ_G0019675 | MGP_PWKPhJ_G0019422 | MGP_PWKPhJ_G0019671 | |
| *Cast* | MGP_CASTEiJ_G0019922 | MGP_CASTEiJ_G0019668 | MGP_CASTEiJ_G0019917 | |
| *Dom* | MGP_WSBEiJ_G0019983 | MGP_WSBEiJ_G0019728 | MGP_WSBEiJ_G0019979 | MGP_WSBEiJ_G0019980 |
| *Spret* | MGP_SPRETEiJ_G0019504 | MGP_SPRETEiJ_G0019238 | MGP_SPRETEiJ_G0019499 | |
| *Car* | MGP_CAROLIEiJ_G0018624 | MGP_CAROLIEiJ_G0018369 | MGP_CAROLIEiJ_G0018621 | |
| *Pah* | XM_021179921.1 | XM_021180719.1 | XM_021215385.1 (*Tpbpa*-like)<br>% idenity to *Dom-Tpbpa*: 84.3%<br>% identity to *Dom-Tpbpb*: 90.6% | XM_021216022.1 (*Tpbpa*-like)<br>% idenity to *Dom-Tpbpa*: 92.2%<br>% identity to *Dom-Tpbpb*: 82.7% |

Ensembl gene stable ID of included sequences for *Mus m. castaneus (Cast), Mus m. musculus (Musc), Mus m. domesticus (Dom), Mus spretus (Spret), Mus caroli (Car)*. NCBI sequence ID of included sequences for *Mus pahari (Pah).*
